# The urinary pathobiont *Actinobaculum massiliense* generates androgens via the *dirAB* pathway

**DOI:** 10.64898/2025.12.18.695155

**Authors:** Taojun Wang, Saeed Ahmad, Raissa Santos de Lima Rosa, Briawna Binion, Francelys V. Fernandez-Materan, Jimoh Olamilekan Igbalaye, David Chung, Anika Bushra, Vince Perez, Michael A. Biedak, Elizabeth Tang, Betsy Barnick, Damie Olukoya, Pauline Mbuvi, Debapriya Dutta, John W. Erdman, H. Rex Gaskins, Glen Yang, Joseph Irudayaraj, Rafael C. Bernardi, Jason M. Ridlon

**Affiliations:** Carl R. Woese Institute for Genomic Biology, Urbana, IL 61801 USA; Department of Animal Sciences, University of Illinois at Urbana-Champaign, Urbana, IL 61801 USA; Department of Bioengineering, University of Illinois at Urbana-Champaign, Urbana, IL 61801 USA; Department of Physics, Auburn University, Auburn, AL, USA; Department of Chemistry and Biochemistry, Auburn University, Auburn, AL 36849, USA; Department of Physics, University of Illinois Urbana-Champaign, Urbana, IL, USA; Biomedical Research Center, Mills Breast Cancer Institute, Carle Foundation Hospital, Urbana, IL, USA; Department of Urology, Carle Foundation Hospital, Urbana, IL, USA; Cancer Center at Illinois, University of Illinois at Urbana-Champaign, Urbana, IL 61801 USA; Division of Nutritional Sciences, University of Illinois at Urbana-Champaign, Urbana, IL 61801 USA; Carle-Illinois College of Medicine, University of Illinois Urbana-Champaign, Urbana, IL, USA; Center for Advanced Study, University of Illinois Urbana-Champaign, Urbana, IL, USA; Department of Microbiology and Immunology, Virginia Commonwealth University School of Medicine, Richmond, VA, USA

## Abstract

While overlooked during the Human Microbiome Project, characterizing the urinary microbiota in health and disease is a new frontier in microbiome science. Recent studies have associated differential abundance of bacterial taxa including *Propionimicrobium lymphophilum* and *Actinobaculum*/*Actinotignum* spp. with prostate cancer. In this study, we collected urine from subjects prior to prostate biopsy and applied a novel <u>H</u>uman <u>S</u>terolbiome <u>D</u>iscovery <u>H</u>igh-throughput (HSDH) assay to identify culturable urinary bacteria with the ability to generate androgens. Application of the HSDH assay to urine samples led to the isolation of eight *P. lymphophilum* strains positive for cortisol side-chain cleavage (steroid-17,20-desmolase), 17β-HSDH activity, or both. In addition, we isolated three strains of *Actinobaculum massiliense* that encode *<u>D</u>HEA isomerase reductase (dir*) genes. The *dirA* gene encodes a novel 3β/17β-hydroxysteroid dehydrogenase/Δ^4,5^-isomerase and the *dirB* gene encodes a novel 17β-hydroxysteroid dehydrogenase isoform. Structural prediction and molecular dynamics reveal probable catalytic mechanisms based on the shared catalytic triad but distinct binding pocket geometries of the DirA and DirB that describe their respective reactions. Phylogenetic analysis of DirA and DirB revealed homologs in urinary tract commensals as well as bacteria associated with steroid degradation found in aquatic and terrestrial environments. Taken together, the development of the HSDH assay and the identification of the *dir* pathway genes is a significant advance in microbial endocrinology, laying the methodological foundation and providing the molecular basis for understanding the role of urinary tract bacteria in host endocrine physiology.

## Introduction

There is a renaissance in interest in the gut microbiome, bile acid, and steroid hormone metabolism^1–4^ that has emerged in the post-genomic era, following a solid foundation that was built during the final decades of the 20th century^5^. Recent studies have reported novel microbial enzymatic functions involved in the biotransformation of steroid hormones and bile acids in the gastrointestinal (GI) tract including dehydroxylation^3,6^, conjugation with sulfate and amino acids^7^, oxidation and epimerization of steroid hormone and bile acid A/B rings^8,9^, and oxidation and epimerization of hydroxyl groups^10–12^. Studies that have measured serum, intestinal, and/or organ steroid hormone and bile acid profiles between conventional and germ-free^13,14^ and broad spectrum antibiotic treatment^15^ demonstrate that commensal gut bacteria significantly contribute to the host steroid hormone and bile acid metabolome. Many of these reactions, such as “toggling” between A/B ring and hydroxyl conformations, are indistinguishable from host metabolic capabilities; whereas reactions, such as 7-dehydroxylation of bile acids^16^ formation of non-canonical amino acid-bile acid conjugates^17^, and the side-chain cleavage of glucocorticoids to 11-oxy-androgens^18^, are thought to be uniquely microbial reactions.

Beyond the GI tract, steroids are concentrated in other body habitats that are also teaming with microbes including the skin, oral cavity, and urinary tract^19^. Until recently, urine was thought to be sterile, and, indeed, characterizing the urinary microbiome was not included as part of the Human Microbiome Project^20^. Instead, pioneering work by researchers studying the urogenital tract and its disorders and cancers has revealed a microbial community distinct from skin and fecal microbiomes^21–25^. This work has largely focused on comparing relative abundances of urinary microbiota between healthy controls and disease states, and has identified differential abundances of certain taxa with diseases, particularly prostate cancer. These taxa include genera, such as *Cutibacterium, Propionimicrobium, Peptoniphilus,* and *Actinobaculum*, elevated in the urine of prostate cancer patients^21,23–25^. Whether these observations are a result of cause or consequence of the changing environment created by prostate tumors is currently unclear. Little is known about the biology, particularly the metabolism of host urinary components by these bacteria, that might contribute to disease processes.

In this regard, our prior work identified a multi-gene, multi-species pathway involved in the microbial conversion of C_21_ cortisol derivatives to derivatives of the androgen-precursor C_19_ 11β-hydroxyandrostenedione (11OHAD) by the gut bacteria, *Clostridium scindens* and *Butyricicoccus desmolans*^18,26,27^. To our surprise, when we performed phylogenetic analysis of the *desE* gene encoding steroid 20β-hydroxysteroid dehydrogenase (20β-HSDH)^28^, and the steroid-17,20-desmolase encoded by the *desA* and *desB* genes^27^ we determined that urinary tract bacteria, such as *Propionimicrobium lymphophilum* harbored the *desABE* genes^27^. Subsequently, we cultured bacteria from urine samples and prostatectomy tissue collected from patients diagnosed with prostate cancer and age-matched control men, and identified bacterial strains that generated androgens from cortisol and/or 11OHAD ^1^. We identified a gene encoding NADPH-dependent 17β-HSDH in the urinary tract commensal, *Propionimicrobium lymphophilum*, which we named *desG*^1^. Importantly, oxygen-independent bacterial desmolase (steroid transketolase encoded by the *desA* and *desB* genes) is expected to catalyze side-chain cleavage of steroids through a mechanism distinct from oxygen-dependent host desmolase (cytochrome P450 monooxygenase)^18,29^. As prostate cancer is a disease driven by androgens, the drugs abiraterone acetate, which inhibits the key steroidogenic cytochrome P450 monooxygenase CYP17A1, and the replacement glucocorticoid, prednisone, are prescribed to block androgen production in advanced prostate cancer. Importantly, we demonstrated that abiraterone does not inhibit gut or urinary bacterial desmolase conversion of cortisol (or prednisone) to androgens, and that prednisone is converted to an androgen that drives prostate cancer cell proliferation^1^. Thus, from a clinical perspective, the human microbiome is expected to be a source of androgens that may facilitate prostate cancer progression.

These studies lay the groundwork for envisioning a potential causal role for the urinary microbiome in the progression of prostate cancer through production of potent androgens (e.g. T, dihydrotestosterone (DHT), 11keto-T, Δ^1,4^-11keto-AD (AT)) from inactive precursors (e.g. cortisol, prednisone, dehydroepiandrosterone (DHEA), androstenedione, 11OHAD) that represent a currently unrecognized environmental source of androgens in the prostate^30^. Despite these recent advances, the diversity of urinary microbial taxa that express 17β-HSDH is currently unknown. Searching current sequence databases with DesG as a query identifies proteins with high amino acid sequence identity, yet fails to take into account that different gene families (i.e. short chain dehydrogenase/reductase (SDR), aldo-keto reductase (AKR), medium chain dehydrogenase (MDR)) have all been reported to encode distinct isoforms of HSDH enzymes, including 17β-HSDH^31^. Taxonomic profiling that relies on 16S rRNA sequencing is useful in determining community composition but requires prior knowledge of species and strain steroidogenesis. Shotgun metagenomic sequencing of a urine sample would provide both taxonomic information and gene content; however, the SDR, AKR, and MDR are very large protein families with diverse substrate preferences (e.g. sugars, alcohols, dyes, sterols) making bioinformatic prediction of steroidogenic sequences daunting without bona fide sequences of high amino acid sequence identity available. AI-based predictions have been able to utilize active-site geometry to identify novel bile acid metabolizing enzymes in the metagenome, but require prior experimental structural data from steroidogenic enzymes to train the model^32^.

Here, we have developed a functional enzyme-coupled assay, <u>H</u>uman <u>S</u>terolbiome <u>D</u>iscovery <u>H</u>igh-throughput (HSDH) assay, to rapidly screen culturomic and/or functional metagenomic libraries for 17β-HSDH activity in a species and sequence-independent and agnostic manner. Moreover, we identified a urinary species that was previously not known to metabolize steroids which was detected by the HSDH assay to express 17β-HSDH activity. One of these isolates, *Actinobaculum massiliense* strain P19, was determined to convert the testicular and adrenal androgen precursor, DHEA, to androstenedione/androstenediol and testosterone. To further mechanistically investigate androgen biosynthesis, genes encoding a novel multi-functional 3β-HSDH-Δ^4–5^-isomerase/17β-HSDH enzyme and a separate 17β-HSDH were identified, and the structures were predicted along with ligand binding and catalysis using advanced AI and molecular dynamics. These findings substantially expand our knowledge of steroid microbiology and lay the groundwork for understanding the role of urinary tract bacteria in host endocrine physiology.

## Results

### Development of the HSDH enzyme-coupled assay

Screening bacterial bioconversion using chromatographic methods has been a standard means to identify steroid-metabolizing bacteria for over 60 years^16,33^. This approach is laborious, time-consuming, and expensive, requiring expertise and access to chromatography instruments (GC/HPLC or GC-MS/LC-MS) to determine which bacterial colonies metabolized steroids. Indeed, our recent study utilized an LC-MS platform to screen urinary cultures for the side-chain cleavage of cortisol and 17-keto reduction^1^. In the search for an alternative approach to screen cultures, we developed a modular assay that utilizes stereo- and regio-specific enzymes, known as hydroxysteroid dehydrogenases (HSDHs), that allow steroid substrates or products to be detected by measuring the change in absorption of cofactors during enzymatic reactions in a plate reader. This HSDH Assay starts with incubation of colonies in a growth medium containing steroid substrates (**Fig. 1a**). We then extract the steroid metabolites from the medium for each isolate with organic solvent, evaporate the solvent, and add a buffer containing NAD(P)H/NAD(P)^+^ and HSDH enzyme to initiate the reaction (**Fig. 1a**). In the current process, bacterial colonies of interest will convert cortisol to 17-keto steroids (e.g. 11OHAD) or convert 11OHAD to 11β-hydroxy-testosterone (11OHT) (**Fig. 1b**). The 96-well plate is then immediately read on a plate reader at 340 nm. Since cortisol is not a substrate for 17β-HSDH, wells in which bacteria did not convert cortisol to a 17keto- product will not yield a ΔAbs _340nm_. If, however, a bacterial colony converts cortisol to 11β-OHAD (17-keto), we will detect a decrease in Abs_340 nm_ according to the reaction:

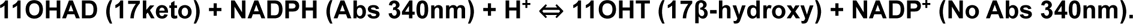

**Figure 1:**
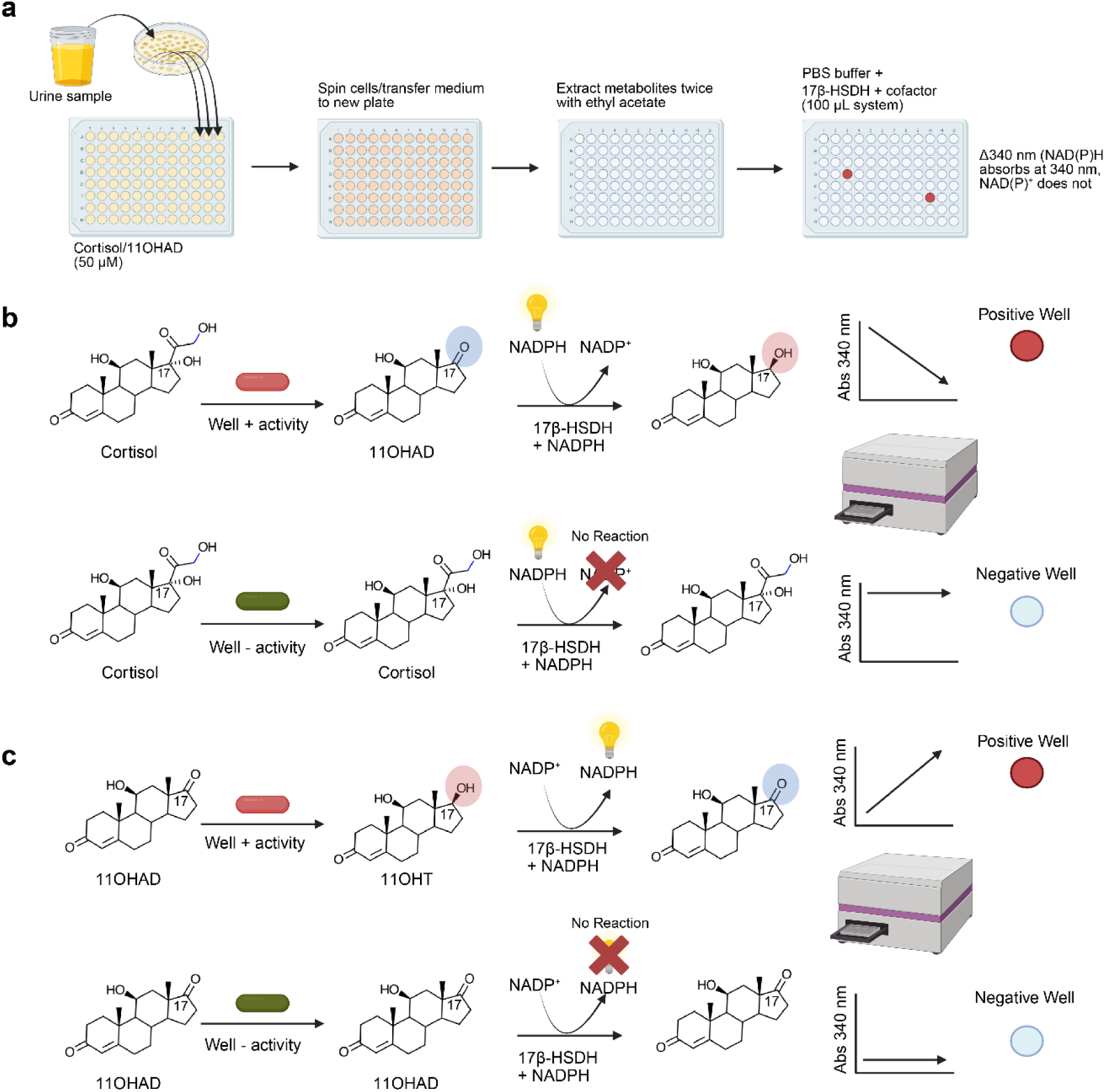
Schematic overview of the Human Sterolbiome Discovery High-throughput (HSDH) assays for urinary bacterial isolation using cortisol or 11OHAD as substrates. **a**, A workflow to prepare the samples for the metabolite analysis with the HSDH assays. Cortisol or 11OHAD was used as a substrate for bacterial isolation. The metabolites were extracted and used for the analysis with 100 uL phosphate-buffered saline (PBS) containing 500 nM 17β-HSDH and 100 µM cofactors (NADPH or NADP^+^). **b**, Bacterial isolates that have the genes encoding steroid-17, 20-desmolase (DesAB) convert cortisol to 11OHAD, subsequently, the added 17β-HSDH converted 11OHAD to 11OHT, concomitantly NADPH was converted to NADP^+^, causing the decrease of the optimal density overtime at 340 nm. While the isolates that do not have the *desAB* genes cannot metabolize cortisol, no change of optimal density can be observed. **c**, Bacterial isolates that have the gene encoding 17β-HSDH convert 11OHAD to 11OHT, subsequently, the added 17β-HSDH converted 11OHT to 11OHAD, concomitantly NADP^+^ was converted to NADPH, causing the increase of the optimal density overtime at 340 nm. While the isolates that do not have this gene cannot metabolize 11OHAD, no change of optimal density can be observed.

Some bacteria catalyze an additional reaction in which the resulting 17keto-steroid is reduced to 17β-hydroxy, which would also result in a negative reaction on the plate reader when NAD(P)H is the cofactor, despite side-chain cleavage (**Fig. 1c**). By running duplicate plates with 17β-HSDH in the presence of NAD(P)^+^, we detect an increase in ΔAbs _340nm_ in those wells in which a 17β-hydroxy androgen is produced from cortisol. Hence, the HSDH assay allows time and cost-effective screening of bacterial urinary/prostate isolates for androgen formation.

### Validation of the HSDH enzyme-coupled assay

We chose the 17β-HSDH enzyme from the fungus *Cochliobolus lunatus* to initiate the reactions due to its favorable bidirectionality^18^. The *C. lunatus* 17β-HSDH cDNA sequence was cloned into pET-51b(+) plasmid and transformed into *E. coli* BL21-CondonPlus-(DE3)-RIPL for the protein production and purification based on the Strep-Tactin affinity (**Extended Data Fig. 1a**). The recombinant protein converted 11OHAD to 11OHT with concomitant conversion of NADPH to NADP^+^. The recombinant protein is also able to perform the reverse reaction in which 11OHT is converted to 11OHAD with simultaneous conversion of NADP^+^ to NADPH (**Extended Data Fig. 1b-c**). NADPH has absorbance at 340 nm, while NADP^+^ does not. Absorbance changes were observed over time when using 50 µM 11OHAD or 11OHT (**Extended Data Fig. 1d-e**) as substrates with 100 µM cofactors added. Furthermore, we investigated the concentrations of the substrates (11OHAD and 11OHT) and the ratio of the substrates in the HSDH Assay. We found that the low concentrations of the substrates can be detected, and the presence of the metabolite by the recombinant protein did not affect the application of the HSDH Assay (**Extended Data Fig. 2**).

To validate the HSDH Assay, we performed control reactions with *E. coli* MG1655 that is not observed to metabolize cortisol, *P. lymphophilum* ANSC4 that is capable of converting cortisol to 11OHAD, and *P. lymphophilum* API-1 that can convert cortisol to both 11OHAD and 11OHT^1^. Each strain was incubated using 50 µM cortisol as a substrate and LC-MS analysis confirmed the activities of each bacterial strain (**Fig. 2a-b**). Afterwards, the spent culture medium from each strain was extracted with ethyl acetate. The extracts were resuspended with 100 µL PBS buffer, 100 µM cofactors and 500 nM recombinant 17β-HSDH enzyme to initiate the reactions. Consistently, for the extracts from *E. coli* MG1655, no absorbance change was observed in either the reductive or oxidative reactions; for the extracts from *P. lymphophilum* ANSC4, absorbance change in reductive reaction was observed over time; for the extracts from *P. lymphophilum* API-1, absorbance change in both reductive and oxidative reactions was observed over time (**Fig. 2c-e**). These results revealed that HSDH Assay is a reliable method to analyze microbial metabolism activities.

**Figure 2:**
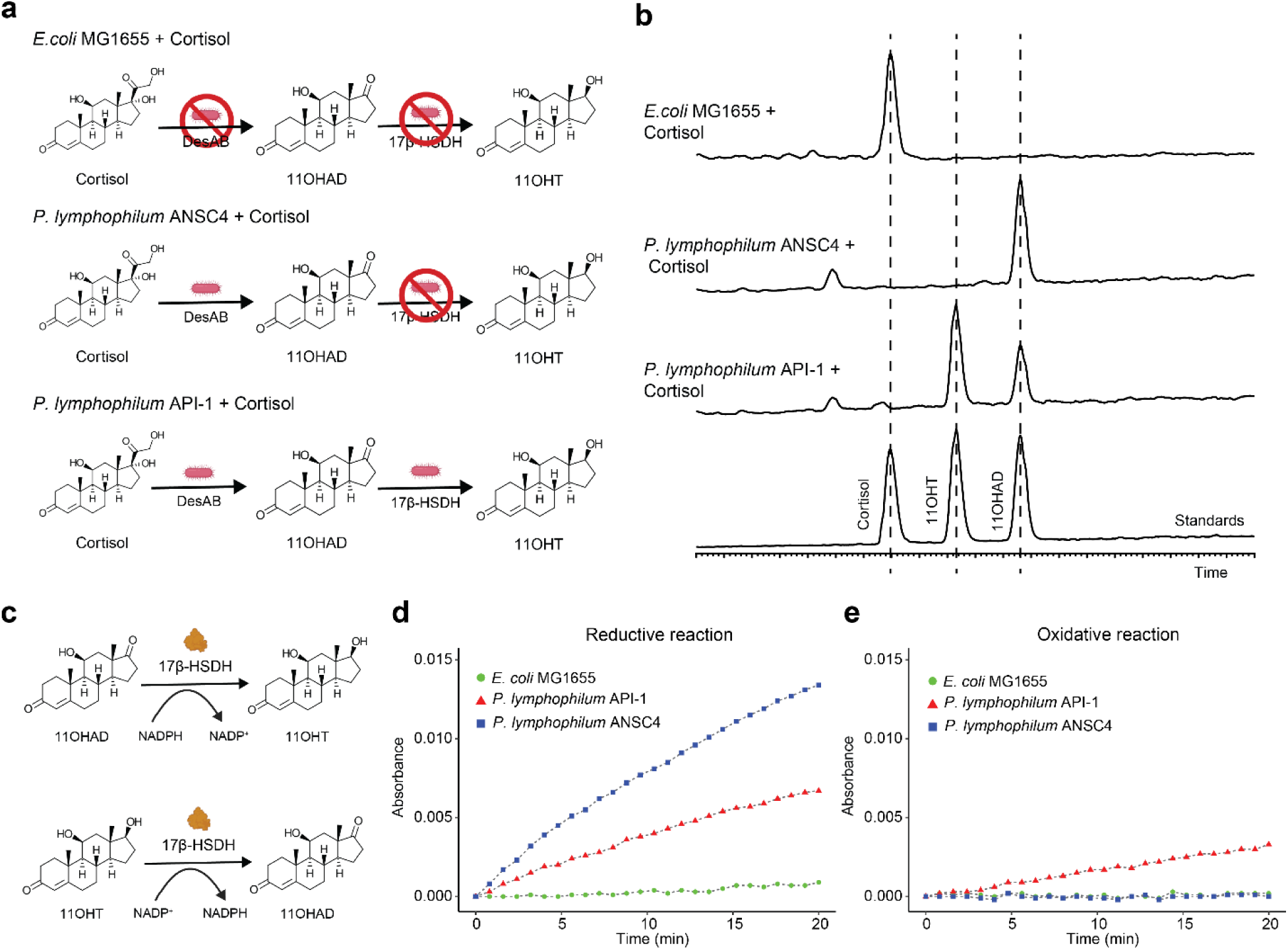
Validation of the HSDH assays. **a,** *E.coli* MG1655 that cannot metabolise cortisol, *P. lymphophilum* ANSC4 that can convert cortisol to 11OHAD and *P. lymphophilum* API-1 that can convert cortisol to 11OHAD and 11OHT were used to validate the HSDH assays. **b,** LC-MS chromatograms showed the metabolites of *E.coli* MG1655, *lymphophilum* ANSC4 and *P. lymphophilum* API-1. **c,** 17β-HSDH from *Cochliobolus lunatus* was used for the HSDH assays. It converted 11OHAD to 11OHT using NADPH as a cofactor, and converted 11OHT to 11OHAD using NADP^+^ as a cofactor. **d,** The change of the optimal density (absorbance) of the three strains using NADPH as the cofactor overtime at 340 nm. **e,** The change of the optimal density (absorbance) of the three strains using the NADP^+^ as cofactor overtime at 340 nm.

### Urine bacterial isolation for steroid metabolism

Building on our previous study^1^, to identify microbial pathways involved in steroid metabolism and androgen production in the male urinary tract, we recruited 27 additional patients at Carle Foundation Hospital for urine collection prior to prostate biopsy (**Supplementary Table 1**). Clean catch mid-stream urine samples were collected prior to biopsy procedure and cultured on several enriched agar media. Resulting bacterial colonies were transferred to 96-well plates to screen for cortisol side-chain cleavage activity or 17β-HSDH using the HSDH assay. To identify wells positive for activity, the medium from each colony inoculated well was extracted after 7 days anaerobic cultivation, evaporated and resuspended in buffer for the HSDH assay. Wells positive for DesAB and/or 17β-HSDH activities were further confirmed by LC-MS analysis (**Extended Data Figure 3**). Subsequent bacterial identification (whole genome sequencing) showed the presence of *Propionimicrobium lymphophilum* and *Actinobaculum massiliense* strains. Among eight *P. lymphophilum* isolates, five express steroid-17,20-desmolase and 17β-HSDH activities, two express steroid-17,20-desmolase activity only and notably one harbored only 17β-HSDH activity. Additionally, three *A. massiliensis* strains were found to express 17β-HSDH activity (**Table 1**). Here 9 out of 27 patients were positive for steroid-17,20-desmolase and/or 17β-HSDH encoding bacteria, where 7 were diagnosed with prostate cancer. All three *A. massiliense* strains were isolated from 3 male patients who were subsequently diagnosed with stage pT1C (CFH51), and T3B (CFH19, CFH34) (**Table 1**; **Supplementary Table 1**).

**Table 1:** Androgen producing bacteria isolated from urine.

| Microbial isolates | Steroid-17,20-desmolase (DesAB) | 17β-Hydroxysteroid dehydrogenase (17β-HSDH) | 3β-HSD/$\Delta^{4,5}$-isomerase |
| --- | --- | --- | --- |
| *A. massiliense* CFH19 |  | + | + |
| *P. lymphophilum* C741 | + | + |  |
| *A. massiliense* CFH34 |  | + | + |
| *P. lymphophilum* P357 | + | + |  |
| *P. lymphophilum* P381 | + | + |  |
| *P. lymphophilum* P4421 | + |  |  |
| *P. lymphophilum* P4435 | + | + |  |
| *P. lymphophilum* P4841 | + |  |  |
| *P. lymphophilum* P4844 |  | + |  |
| *A. massiliense* CFH51 |  | + | + |
| *P. lymphophilum* P5134 | + | + |  |

### Pangenome analysis of Propionimicrobium lymphophilum strains

Under our culture conditions, *P. lymphophilum* is the most isolated steroid metabolizing and sole glucocorticoid side-chain cleaving taxon. To comprehensively investigate this species, 22 *P. lymphophilum* strains were included for pangenome analysis. Genomic characteristics of the strains are shown in (**Supplementary Table 2-3)**. Both BUSCO and checkM were used to assess the completeness of the genome assembly, and all of them showed a high completeness and low contamination (**Supplementary Fig. 1**). The Roary analysis identified a pangenome containing 4565 genes with 1422 in the core genes (**Fig. 3a**). The gene function was distributed in information storage, cellular processes, metabolism or poorly characterized (**Supplementary Fig. 2, Supplementary Table 4**). The gene number of each strain ranged from 1865 to 2192 (**Fig. 3b, Supplementary Table 3**). The type strain DSM 4903, isolated from human submaxillary tissue, has the highest number of unique genes with a total of 156 genes, followed by API-1 with 129 genes isolated from homo sapiens prostate tissue. Pan-core plot (**Fig. 3c**) showed the relationship between the pangenome, core genome and the number of genomes. The number of genes in the pangenome increased, and the core genes in the core genome decreased with each consecutive addition of a *P. lymphophilum* genome. Heap’s Law analysis showed that the alpha value is 1.00, indicating a closed pangenome.

**Figure 3:**
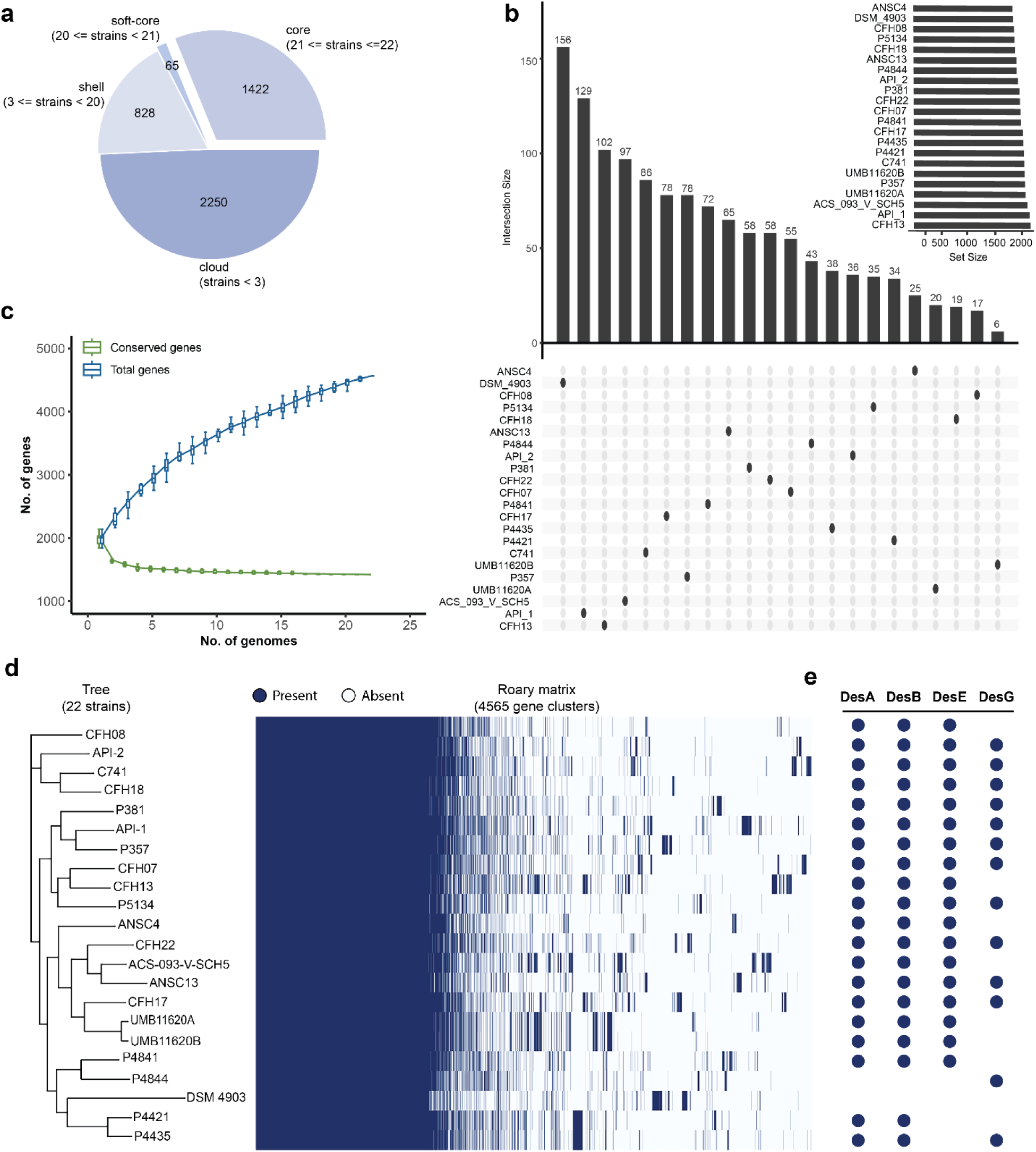
Pangenome analysis of *Propionimicrobium lymphophilum* strains. **a,** The size of the core genes, shell genes, and cloud genes. **b,** The gene size and the unique gene of each *P. lymphophilum* strain. **c,** Pan-core plot for the 22 *P. lymphophilum* strains. **d,** Phylogenetic tree and matrix of presence and absence of the core and accessory genes of the pangenome. **e,** The presence and absence of the genes that are associated with steroid metabolism.

The average nucleotide identity analysis (**Extended Data Fig. 4**) showed that all the strains share > 98% identity, indicating all of them have very similar genomes. To further compare the gene composition between strains, a phylogenomic tree was created based on the presence and absence of the genes (**Fig. 3d**). DesA, DesB, DesE, and DesG amino acid sequences were searched among all the genomes, and their presence and absence showed diversity among strains for steroid metabolic capacity (**Fig. 3e**). Interestingly, 12 of the *P. lymphophilum* strains from urine have both the *desAB* and *desG* genes; 8 strains only have *desAB* genes, and 1 strain only has *desG* gene. The genome of the type strain DSM 4903 lacks *desAB* and *desG* genes. To further confirm this, strain DSM 4903 was cultivated in the presence of cortisol or 11OHAD as substrate. Consistent with the genome sequence, we found that steroid substrates were not metabolized by strain DSM 4903 (**Extended Data Fig. 5**). Moreover, we purchased another commercially available ACS-093-V-SVH5 strain that contains *desAB* genes, and incubated it with cortisol or 11OHAD as substrate, finding that 11OHAD was produced while 11OHT was not produced (**Extended Data Fig. 5**).

KEGG analysis of the core genome of *P. lymphophilum* suggests that this bacterium is a strictly anaerobic, peptide-fermenting organism that relies on simple and highly efficient metabolic routes to survive urinary or mucosal niches (**Supplementary Table 5**). It encodes a complete Embden-Meyerhof-Parnas pathway and a broad non-oxidative pentose phosphate pathway. However, the Entner-Doudoroff pathway is mostly missing, and the TCA cycle is shown as open due to lacking the isocitrate dehydrogenase step, found to be a common feature on gram-positive anaerobic bacteria^34^. KEGG analysis also show that *P. lymphophilum* carries multiple transport systems for peptides, metals, and nutrients such as Opp, ZnuABC, MetQIN, and CbiNMQO, suggesting some dependency on host derived substrates, a behavior commonly observed in host-associated opportunistic pathogens in nutrient limited environments and likely metabolic flexibility^35^. Even though some amino acid biosynthesis pathways seem to be incomplete, like asparagine, arginine, tyrosine, phenylalanine, and tryptophan pathways, many vitamin and cofactor pathways, including folate/THF, pyridoxal-5′-phosphate, riboflavin derived FMN/FAD, and lipoic acid, are largely present, suggesting that this bacterium can maintain redox balance even when regular carbon sources are scarce^36^.

### *Actinobaculum massiliense* converts DHEA to testosterone

Three strains of *A. massiliense* isolated from human urine samples express 17β-HSDH activity (**Table 1**). We obtained a scanning electron micrograph of *A. massiliense* CFH19 (**Fig. 4a**) confirming the curved coccobacillus morphology, as well as evident extracellular polysaccharides connecting microbial cells. We obtained a circular 2.03 Mbp genome of *A. massiliense* CFH19 and observed a gene annotated as “3β-HSD/Δ^4,5^-isomerase” suggesting metabolism of the adrenal androgen precursor DHEA to androstenedione (**Fig. 4b**). Thus, we propose that *A. massiliense* CFH19 can convert DHEA to androstenedione/androstenediol to testosterone (**Fig. 4c**). To test this hypothesis, we then confirmed the metabolism of DHEA, observing the consistent decrease of DHEA from 24 hr (26.25 ± 0.70 µM) to 48 hr (11.02 ± 0.04 µM) and 72 hr (5.79 ± 0.27 µM) (**Fig. 4d**). Concomitantly, there was a rise in androstenedione at 24 hr (15.29 ± 0.38 µM) followed by consumption of androstenedione at 48 hr (8.12 ± 0.51 µM) and 72 hr (4.36 ± 0.34 µM). From 24 hr we measured production of androstenediol (5.74 ± 0.36 µM) and testosterone (4.14 ± 0.07 µM), which rose in concentration at 48 hrs (16.90 ± 1.56 µM, 12.48 ± 0.47 µM, respectively) and 72 hrs (21.45 ± 1.76 µM, 17.15 ± 0.45 µM, respectively) (**Fig. 4d**). In humans, DHEA can be converted to testosterone via two ways: 1), DHEA to androstenedione via HSD3B1 and HSD3B2 activities, then androstenedione to testosterone via 17HSDH activity; 2), DHEA to androstenediol via 17HSDH activities, then androstenediol to testosterone via HSD3B1 and HSD3B2 activities^37^. To further investigate which intermediates are used for testosterone biosynthesis, we tested androstenedione and androstenediol as substrates. *A. massiliense* CFH19 was capable of converting both substrates to testosterone. The addition of androstenedione resulted in formation of DHEA, androstenediol, and testosterone during incubations (**Fig. 4e**). Similarly, the addition of androstenediol yielded DHEA, androstenedione, and testosterone during incubations (**Fig. 4f**). Therefore, we propose that DHEA can be converted to androstenedione or androstenediol and from each to testosterone in *A. massiliense* CFH19.

**Figure 4:**
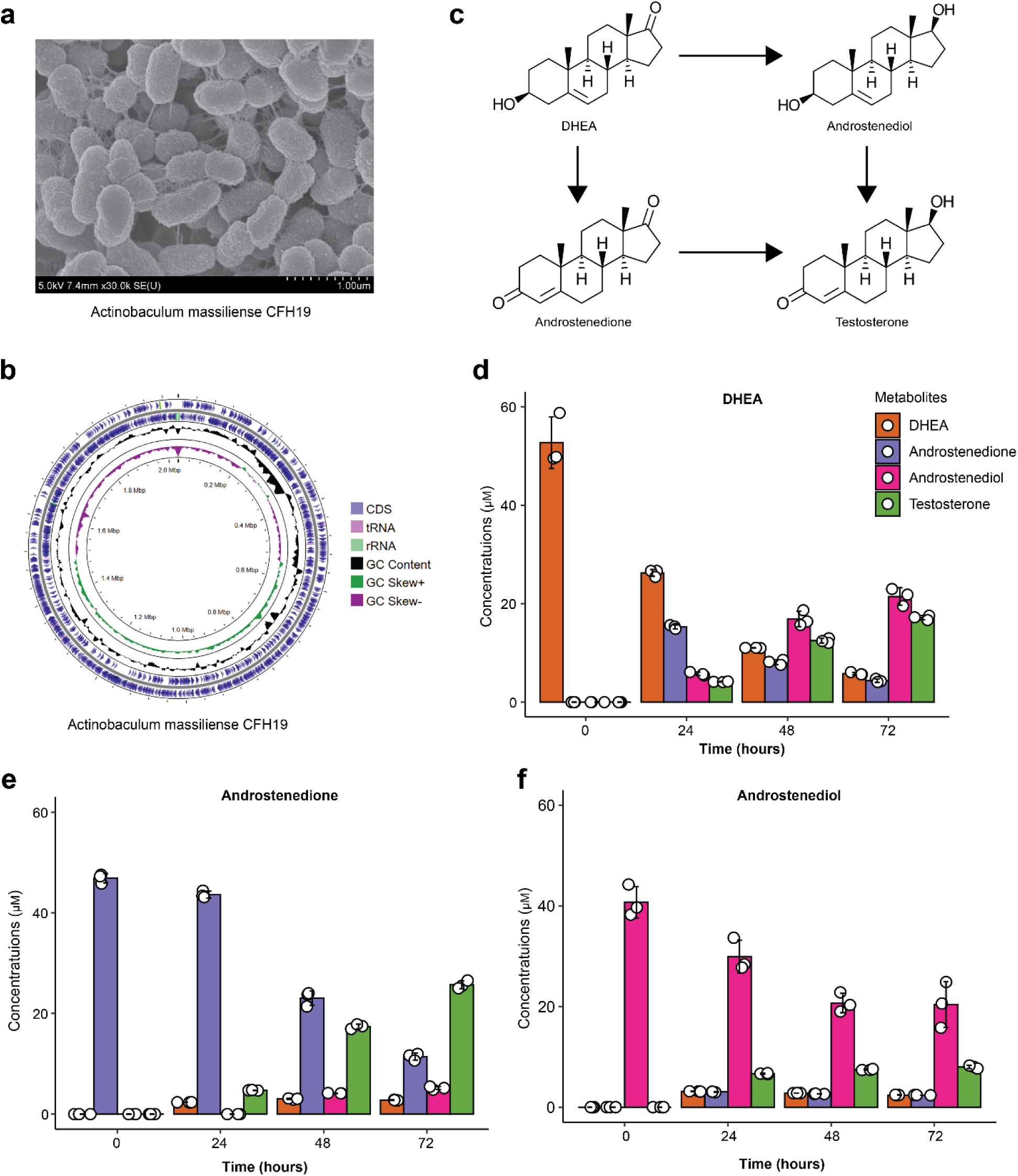
Androgen biosynthesis by *Actinobaculum massiliense* CFH19. **a,** SEM image of *A. massiliense* CFH19. **b,** The circular genome map of *A. massiliense* CFH19. **c,** The proposed bacterial pathways for testosterone production using DHEA as substrate. *A. massiliense* CFH19 metabolises DHEA (**d**), androstenedione (**e**) and androstenediol (**f**) to produce testosterone over time.

### Identification of genes encoding 3β/17β-HSDH/Δ^4,5^-isomerase and 17β-HSDH

To further investigate which genes are involved in the bacterial enzymatic pathways, we examined the genome for annotations consistent with protein families known to have 17β-HSDH activity (e.g. short chain dehydrogenase/reductase, medium chain reductase, and aldo-keto reductase). We found two genes encoding putative “3ɑ-(or 20β)-hydroxysteroid dehydrogenase” (PKOOAGGE_00485) and “3ɑ-hydroxycholanate dehydrogenase” (PKOOAGGE_00635) (**Supplementary Table 6**). We then had the PKOOAGGE_00485 (gene 485) and PKOOAGGE_00635 (gene 635) gene candidates synthesized in the codon usage of *E. coli* BL21-CodonPlus(DE3)-RIPL for ectopic expression from the pETIDT-C-His plasmid (**Fig. 5a**).

**Figure 5:**
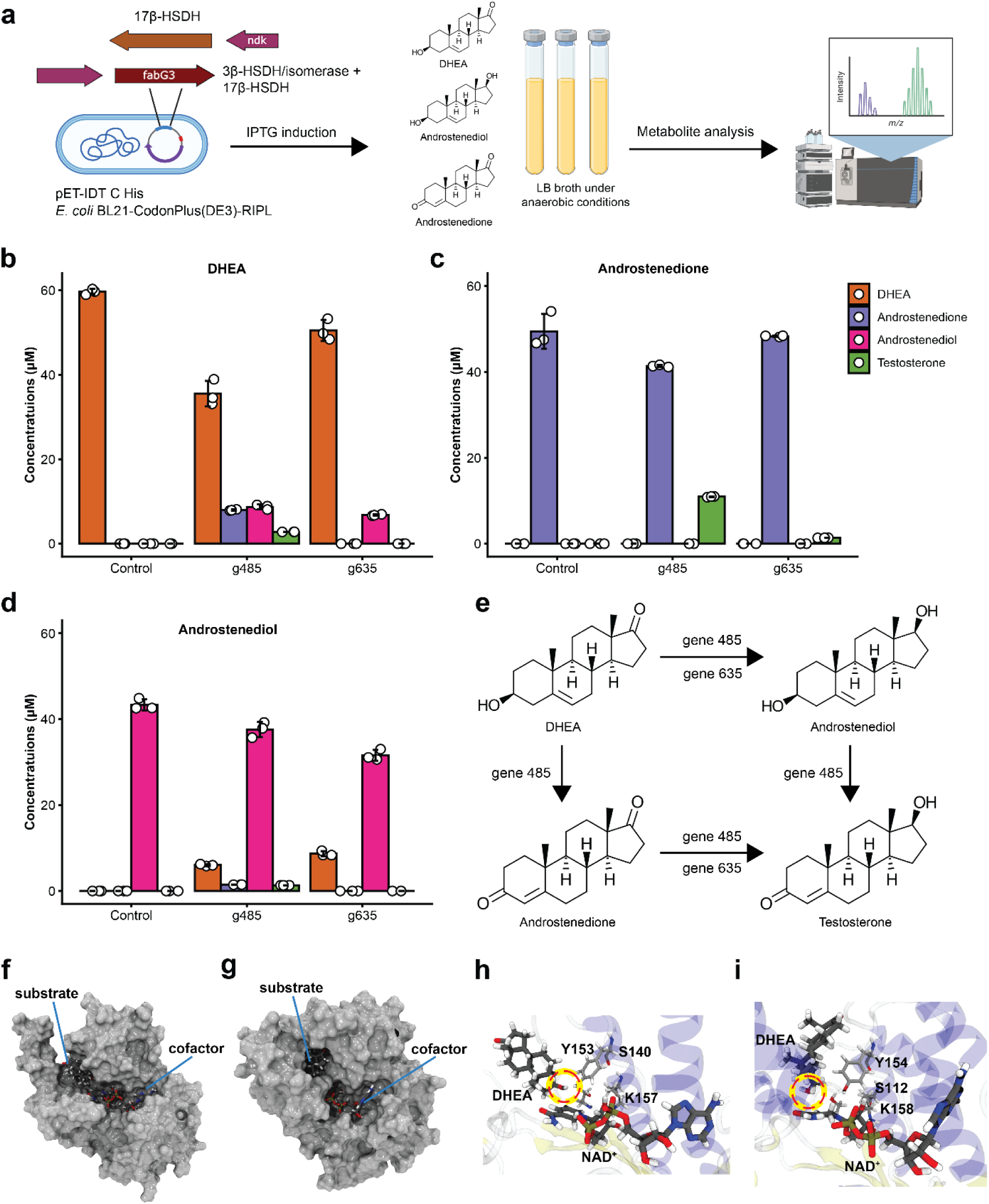
Identification of the genes involved in the androgen biosynthesis. **a,** Two gene predicted to be 3β-HSDH/isomerase and 17β-HSDH were synthesised and transferred into *E. coli* BL21-CodonPlus(DE3)-RIPL) The substrates, DHEA, androstenedione and androstenediol were used to incubate the *E. coli* BL21-CodonPlus(DE3)-RIPL with target genes. Metabolites were analysed with LC-MS. Quantification analysis of the metabolites using DHEA (**b**), androstenedione (**c**) or androstenediol (**d**) as substrates. **e,** The gene functions in the bacterial enzymatic pathways for the androgen biosynthesis. Surface representation of substrate binding to enzymes encoded by gene 485 **(f)** and gene 635 **(g).** Monomeric structures were predicted using AlphaFold3 and shown as molecular surfaces. Substrate molecules and cofactors are displayed in stick representation within the proposed binding pockets. Molecular dynamic trajectory analysis of DirA (**h**) and DirB (**i**) revealed strong interactions between ligands and the catalytic triad.

The expression of gene 485 (SDR family) in *E. coli* (g485) under anaerobic conditions resulted in conversion of DHEA to androstenedione (3β-HSD/Δ^4,5^-isomerase) and androstenedione to testosterone (17β-HSDH) or DHEA to androstenediol (17β-HSDH) and androstenediol to testosterone (3β-HSD/Δ^4,5^-isomerase) (**Fig. 5b-d**). The g485 incubated in the presence of DHEA yielded 7.97 ± 0.14 µM androstenedione, 8.73 ± 0.57 µM androstenediol, and 2.81 ± 0.01 µM testosterone (**Fig 5b**). When androstenedione was added as the substrate, g485 yielded 10.94 ± 0.10 µM testosterone (**Fig 5c**). Addition of androstenediol yielded 1.47 ± 0.01 µM androstenedione, 1.32 ± 0.01 µM testosterone, and 6.01 ± 0.22 µM DHEA. Thus, gene 485 encodes a novel 3β/17β-HSD/Δ^4,5^-isomerase. We suggest the name *<u>D</u>HEA isomerase reductase (dirA*) for gene 485. This is the only 3β-HSD/Δ^4,5^-isomerase that also has 17β-HSDH activity described to date in bacteria, but shares functional similarities with human HSD17B2 (**Fig. 5e**)^38^.

Gene 635 was overexpressed anaerobically in *E. coli* in the presence of 50 µM steroids. When DHEA was added, steroids measured after 24 hrs were DHEA at 50.50 ± 2.51 µM and androstenediol at 6.78 ± 0.18 µM. Androstenedione and testosterone were not detected. Similarly, addition of androstenedione resulted in detection of 48.33 ± 0.16 µM androstenedione and 1.37 ± 0.01 µM testosterone while DHEA and androstenediol were not detected. Addition of androstenediol resulted in 8.69 ± 0.59 µM DHEA, with no detection of androstenedione or testosterone. Taken together, gene 635 encodes a steroid 17β-HSDH that lacks 3β-HSD/Δ^4,5^-isomerase activity. We propose the name *dirB* for gene 635.

### Structural prediction and phylogeny

High-confidence structural models were generated for the enzymes encoded by genes *dirA* and *dirB* using AlphaFold3^39^, with ipTM+pTM values of 0.95 and 0.93, respectively. Both predictions were performed in the presence of NAD⁺ or NADH to stabilize the cofactor-binding region and to ensure that the catalytic clefts were modelled in a catalytically relevant configuration. Although the two enzymes shared an overall conserved fold, a clear structural divergence emerged in the architecture of the substrate-binding pocket. DirA contained a wider and more open catalytic cleft that exposed the catalytic residues Ser140, Tyr153, and Lys157 (**Fig. 5f, h)**. In contrast, the catalytic pocket of DirB was considerably narrower, forming a short and constricted tunnel in which the corresponding residues Ser112, Tyr154, and Lys158 were positioned deeper and with reduced accessibility (**Fig. 5g, i)**. These structural differences suggested that the two enzymes may differ significantly in their ability to orient steroid substrates for catalysis.

Docking studies were consistent with this interpretation. For DirA, DHEA bound favorably in both the 3′ and 17′ orientations that are required for the sequential oxidation and reduction steps, with docking energies of −7.1 and −6.9 kcal mol⁻¹, respectively. Androstenedione and androstenediol also displayed stable and catalytically relevant binding modes, with scores between −6.4 and −6.8 kcal mol⁻¹, respectively. These poses aligned the reactive carbonyl or hydroxyl groups toward the catalytic residues and the NAD⁺ or NADH cofactor in geometries consistent with the proposed reaction sequence (**Fig. 5h**). In contrast, the enzyme encoded by DirB, although capable of binding DHEA with strong affinity in both orientations, often drove the ligand into flipped or misaligned conformations inside the narrower pocket (**Fig. 5i**). Androstenedione bound weakly and consistently failed to orient its 3′ carbonyl in a productive manner. Androstenediol bound tightly but favored a pose that did not support the hydrogen-abstraction chemistry required at the 3′ position.

Using QwikMD^40^ and NAMD3^41^, molecular dynamics simulations further distinguished the catalytic capabilities of the two enzymes. In DirA, all ligand complexes remained stable throughout the 10 ns trajectories. Substrates maintained distances to Ser140 and Tyr153 that were compatible with hydrogen transfer and remained properly oriented toward the NAD⁺ or NADH cofactor (**Fig. 5h**; **Fig. 6**). DHEA in the 3′ orientation remained aligned for oxidative conversion to androstenedione, while androstenedione in the 17′ orientation remained well positioned for reduction to testosterone (**Fig. 6**). DirB, however, consistently failed to maintain such catalytic geometries (**Fig. 5i; Extended Data Figure 6)**. Although substrates remained stably bound inside the tunnel, they did not preserve orientations in which the 3′ carbonyl or hydroxyl groups were accessible to the catalytic residues. Instead, they maintain a flipped non-reactive orientation predicted by the docking simulations. These results support the functional restriction of DirB, which catalyzes only the partial conversion steps DHEA to androstenediol and androstenedione to testosterone, while lacking the structural environment required for 3′-position transformations (**Extended Data Figure 6**).

**Figure 6:**
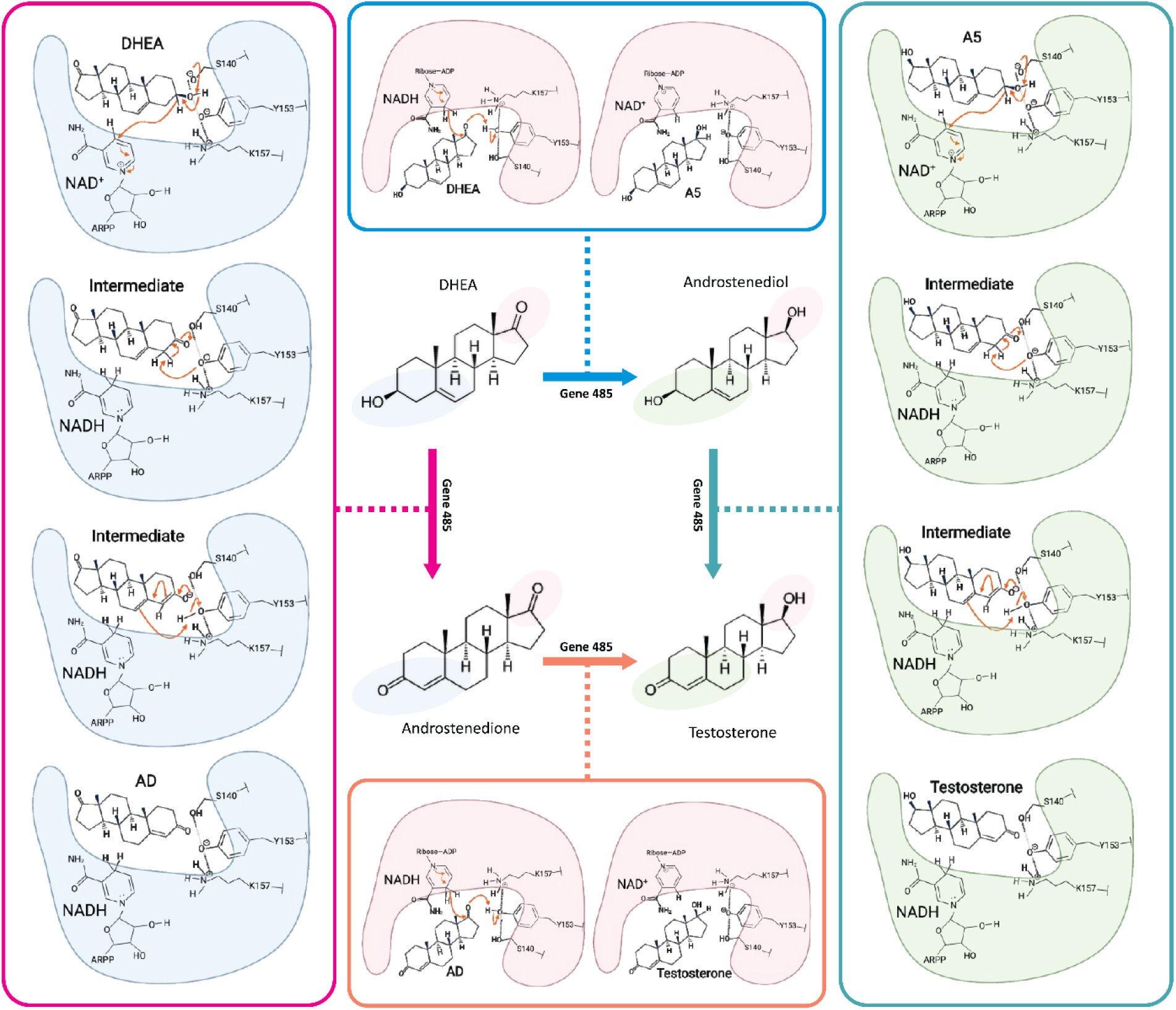
Proposed catalytic mechanism of enzyme DirA (Gene 485) in steroid conversion. Schematic representation of the reaction pathways catalyzed by DirA. The pink pathway (left) illustrates the conversion of DHEA to androstenedione (AD), involving oxidation at the 3′ position via the catalytic triad (Ser140–Tyr153–Lys157). The orange pathway (bottom) depicts the subsequent reduction of androstenedione to testosterone, requiring substrate reorientation (17′ side facing the active pocket) and proton donation by NADH. The blue pathway (top) represents the conversion of DHEA to androstenediol (A5), while the cyan pathway (right) shows the final androstenediol to testosterone step.

### Phylogenetic analysis

To determine the potential species distributions of DirA and DirB, we performed maximum likelihood phylogenetic analysis (**Extended Data Figs. 7 & 8**). The phylogeny of DirA resulted in homologous sequences from multiple phyla including *Bacillota*, *Actinomycetota*, *Pseudomonadota*, *Myxococcota*, *Bacteroidota*, and *Planctomycetota* indicating species distribution in both host-associated as well as aquatic and terrestrial environments. DirA clustered with other sequences from *Actinomycetota* which included sequences from *Corynebacterium* and *Nocardioides. Nocardioides simplex* harbors dozens of genes involved in aerobic sterol degradation, including the *ksi* gene encoding 3β-HSDH/Δ^4,5^-isomerase^42^. The *Nocardioides* sp. WS12 WP_182379233.1 (26.8 kDa) protein shares 27.8% amino acid ID with DirA (28.8 kDa) and encodes a distinct 3β-HSDH/Δ^4,5^-isomerase from *ksi* (13.3 kDa)*. Corynebacterium* spp. are commonly isolated urinary commensals^43^, with some strains isolated from axillary skin that were previously shown to metabolize testosterone, including strains with 3β/17β-HSDH activity^44,45^. More distantly related are proteins from *Nocardia* and *Thermomonospora*, bacteria that have previously been confirmed to metabolize sterols and/or bile acids^46,47^.

Phylogenetic analysis of the DirB gene product revealed homologs in the Actinomycetota, Pseudomonadota phyla. The majority of taxa represented are gram-positive, non-spore forming, aerobic rods common in soil and aquatic and terrestrial environmental samples. DirB homologs were found in numerous *Microbacterium* spp. genomes. Recently, *Microbacterium* spp. were found to be abundant in advanced T3 prostate cancer tumor tissue^48^. These results are expected to expand currently known steroid degrading members of the *Actinomycetota*, and *Pseudomonadota*, and point to novel steroid metabolizing enzymes homologous to DirA and DirB spread across multiple phyla.

## Discussion

The urinary microbiome is emerging as an important factor in urinary tract health and disease, despite not being a part of the Human Microbiome Project^49^. Recent sequenced-based studies of urinary tract microbiome compositions have associated *Propionimicrobium* and *Actinobaculum*^25,50^ with prostate cancer. While cross-sectional studies are not able to determine cause from consequence, or potential mechanisms by which these taxa may potentiate disease development and/or progression, our approach led to identifying androgen-producing taxa that have independently triangulated on these same taxa. Urine is a major route of steroid excretion, composed largely of inactivated androgen precursors (e.g. androstenedione, DHEA, 11OHAD) in levels upwards of several hundred nanomolar in concentration^51^. The reactivation of potent androgens by microbial taxa adhering to urinary epithelial cells, particularly those in the prostatic urethra, provides a proximal route for diffusion of androgens into the prostate (**Extended Data Fig. 9**).

Our recent work established a cortisol side-chain cleavage reaction in gut microbial taxa including *Clostridium scindens*^26^ and *Butyricoccus desmolans*^28^ as well as urinary tract taxa, particularly *P. lymphophilum*^52^. Here, we developed a novel HSDH assay that was used to identify bacterial colonies capable of steroid 17-keto reduction, the assay is modular and can be used to identify steroid biotransformations of glucocorticoids, mineralocorticoids, pregnanes, estrogens, androgens, oxysterols, and bile acids using a myriad of hydroxysteroid dehydrogenase enzymes that provide readouts allowing detection of oxidoreduction, isomerization, lyase, and dehydroxylation reactions when coupled with traditional chromatographic and spectroscopic methods for confirmation. By using this assay, we have extended our culturome to include additional *P. lymphophilum* strains with diverse *des* gene carriage. This culturome is biased in our selection of strains with steroid metabolic function, yet at least one culture collection strain that has been isolated previously was found to lack *des* genes indicating that strain variation will determine the androgenic potential between humans colonized by *P. lymphophilum*. With the available genomes that have been acquired by us and others, we were able to report that the pangenome is now closed, and future work can begin dissecting the biological functions of *P*. *lymphophilum*. In those strains harboring the *desAB* genes, glucocorticoids are converted to 11-oxy-androstanes catalyzed by steroid-17,20-desmolase. The *desC* (20ɑ-HSDH) and *desE* genes (20β-HSDH) function as glucocorticoid side-chain oxidoreductases that regulate recognition by steroid-17,20-desmolase^26,28^. 11-oxy-androstanes are then converted to derivatives of epitestosterone by 17ɑ-HSDH (*desF* gene) or testosterone derivatives by 17β-HSDH (*desG* gene)^1^. The discovery of the *dirA* and *dirB* genes in this study provides another pathway by which commensal urinary tract bacteria generate potent androgen receptor (AR) ligands from weak or non-AR ligand precursors such as DHEA and androstenedione.

In males, the adrenal androgen precursor DHEA is by far the most concentrated urinary steroid with excretion of 1-3 mg per day^51^. Host tissues convert circulating DHEA to androstenedione and testosterone through enzymes with 3β-HSD/Δ^4–5^-isomerase (HSD3B1, HSD3B2) and 17β-HSDH activities (HSD17B3)^53^. The only other enzyme reported to catalyze 3β-HSD/Δ^4-5^ isomerase and 17β-HSDH activities is HSD17B2 which shares only 23.1% amino acid identity with DirA^38^. The current work uncovers a previously unknown metabolic function in which urinary tract bacteria convert DHEA to androstenedione/androstenediol and testosterone. Unexpectedly, these metabolic products of DHEA can be generated by a single enzyme encoded by the *dirA* gene in the short chain dehydrogenase reductase superfamily. A second gene, *dirB,* encodes an enzyme with 17β-HSDH activity. *Comamonas testosteroni* 3β/17β-HSDH shares 45.9% amino acid identity with DirA from *A. massiliensse.* Recent work identified 3β-HSD/Δ^4-5^-isomerase involved in the conversion of pregnenolone to progesterone by the gut microbiome^54^. Specifically, *Mediterraneibacter faecis* and *Floccifex porci* were reported to harbor gene fusions with 3β-HSD/Δ^4-5^-isomerase and 5β-reductase activity resulting in conversion of pregnenolone to 5β-dihydroprogesterone and epiprogesterone by the gut microbiota^54^. These enzymes are distinct from DirA (SDR family; 28.8 kDa) in both size (127.6 kDa) and protein domain architecture (mycofactocin system FadH/OYE family oxidoreductase and TIM barrel fold). Thus, DirA represents a novel 3β/17β-HSD/Δ^4-5^-isomerase, and along with DirB, a pathway by which urinary tract bacteria convert the adrenal androgen precursor, DHEA, into the androgen, testosterone.

The combination of structural modeling, docking, and molecular dynamics provides a mechanistic explanation for the distinct catalytic outcomes of the enzymes encoded by genes *dir*A and *dir*B. Although both proteins retain residues that resemble a classical catalytic triad, their catalytic pockets differ greatly in geometry and accessibility. DirA contains a broad and open cleft that permits substrates to bind in both orientations necessary for the complete reaction sequence. The MD simulations confirm that once the substrate enters the pocket, it maintains productive distances to Ser140 and Tyr153 and remains aligned with the NAD⁺ or NADH cofactor. These features support a classical steroid dehydrogenase mechanism involving coordinated oxidation, reduction, and isomerization steps. This type of reaction sequence has been widely described in steroid dehydrogenases and is consistent with the full catalytic versatility observed for DirA^55–57^.

DirB, by contrast, is limited by its much narrower substrate-access tunnel, which restricts substrate mobility and prevents ligands from achieving catalytically competent orientations. Although this enzyme retains residues that correspond to the same triad, the pocket architecture imposes a strong geometric constraint on ligand positioning. As a result, only reactions that require less precise orientation, such as the reduction of androstenedione to testosterone, are supported. Reactions that require productive 3′-position alignment do not occur because the substrate cannot maintain the necessary geometry inside the tunnel (flipped by 180 degrees). This provides a structural rationale for the reduced catalytic capacity of gene DirB.

It is important to note that the mechanism observed for DirA closely resembles the classical mechanism described for human steroid dehydrogenases^58,59^. Human enzymes, however, often rely on a catalytic tetrad rather than a triad^59–61^. The presence of an additional residue in the human enzymes improves proton relay and stabilizes transition states during both oxidative and reductive steps. The enzymes studied here follow the same basic chemical logic but operate with a more compact catalytic architecture. Despite this difference, the essential orientation requirements and the dependence on precise ligand alignment with the cofactor are conserved, which reinforces the interpretation that pocket geometry, rather than catalytic residue identity, is the primary determinant of functional divergence in this system.

Finally, commensal steroid biotransformations have implications for urinary measurement of steroids for both medical diagnostics as well as determine doping in athletes. The conversion of glucocorticoids and DHEA into derivatives of testosterone and epitestosterone, the ratio of which is used to determine the use of performance enhancing androgens, may be affected by urethral and bladder microbiota^43^. The study of the urinary microbiome (e.g. “urobiome) is in its infancy^43^. The major focus of the field at present is determining the microbiome composition between “healthy” and diseased states, with function as a future goal^43^. Our approach is to define function with respect to mapping the urinary sterolbiome^19^ allowing future work to determine the potential role of microbial androgen formation in the development and progression of androgen-driven diseases, such as prostate cancer^1,62^.

## Method and Materials

### Bacteria and chemicals

#### Bacteria

*Propionimicrobium lymphophilum* strain API-1 was isolated from prostatectomy tissue. *P. lymphophilum* strains API-2, CFH07, CFH08, CFH13, CFH17, CFH18, CFH22, ANSC4, ANSC13, P357, P381, P4421, P4435, P4841, P4844, P5134, C741 were isolated from urine. *P. lymphophilum* strains DSM 4903 and ACS-093-V-SCH5 were obtained from German Collection of Microorganisms and Cell Cultures GmbH, and the Culture Collection, University of Götesborg, Sweden, respectively. *Actinobaculum massiliense* CFH19, *A. massiliense* CFH34, *A. massiliense* CFH5134 were isolated from urine. *E. coli* K-12 MG1655 was provided by the Imlay Lab (James A. Imlay, Professor of Microbiology, University of Illinois Urbana-Champaign); *E. coli* BL21-CodonPlus (DE3)-RIL Competent Cells were purchased from Agilent (Santa Clara, CA, USA).

#### Chemicals

Chemicals included: cortisol (Sigma); 11β-hydroxy-androstenedione (11OHAD, Steraloids, Newport, RI, USA); 11β-hydroxy-testosterone (11OHT, Steraloids); Dehydroepiandrosterone (DHEA, Steraloids); Androstenedione (Steraloids); Androstenedione-2,2,4,6,6,16,16-D7 (Steraloids); Androstenediol (Steraloids); Testosterone (Steraloids); Testosterone-16,16,17-D3 (Steraloids).

#### Bacterial media preparation

Lysogeny broth (LB, Sigma) was purchased and prepared according to the manufacturer’s instructions. Peptone Yeast Glucose (PYG) broth (modified) was prepared according to the DSMZ protocol. Blood agar base (Sigma) was purchased and prepared according to the instructions, with 6% (v/v) defibrinated sheep blood (Thermo Scientific) added. Schaedler agar (Sigma) was purchased and prepared according to the manufacturer’s instructions. Columbia broth (BBL) was purchased and prepared as instructed, with 15 g/L agar added to solidify the medium.

#### Patient recruitment and urine sample collection

To identify androgen forming urinary microbial taxa, we consented and recruited 27 patients under the IRB 22383 (University of Illinois Urbana-Champaign) and Carle Foundation Hospital 18CCC1757. All the study participants had no history of prostate cancer. The age ranged from 50 to 90 years with a BMI less than 35. Patients were excluded if they had active diabetes; were being treated for sexually transmitted infection or urinary tract infection; were taking antibiotics within the past month; or were being treated for benign prostatic hyperplasia, that is, with drugs such as alfuzosin (Uroxatral), doxazosin (Cardura), tamsulosin (Flomax) and terazosin (Hytrin) or abiraterone/prednisone or similar drugs. Participants were given a urine collection kit and detailed instructions on how to properly collect a ‘clean/sterile catch’. The urine collection kit contained an alcohol swab to clean the urethra and tip of the penis. Participants were directed to catch mid-stream urine in sterile vials. At least 20 ml of urine was collected from participants. The urine samples were processed immediately for culturomics work.

#### Urine culturomics

To isolate androgen producing bacteria, 100 µL urine from each patient was plated on the Blood agar, Columbia agar and Schaedler agar. These plates were incubated anaerobically at 37 °C. After 4–5 days, single colonies were picked using sterilized toothpicks and transferred to 96-well plates containing PYG broth medium. After incubation in an anaerobic chamber for 5 days, these plates were utilized as master plates where 50 uL was transferred to deep well plates containing 800 uL PYG medium with either 100 μM cortisol or 11OHAD as substrates. Cortisol was used to identify side-chain cleavage activity (*desAB*) followed by 17β-HSDH activities in the urine yielding 11OHAD and 11OHT. 11OHAD as substrate was used to confirm 17β-HSDH activity where it gets converted to 11OHT when *desAB*-encoding microbes are lacking in the sample. After a week of incubation, these plates were collected and stored at −80°C for screening by the HSDH assay.

#### Heterologous expression and purification of 17β-HSDH from *Cochliobolus lunatus*

The *C. lunatus* cDNA cloning and expression, and protein purification were as described previously^18^. Briefly, recombinant plasmids were transformed into *E. coli* BL-21 CodonPlus (DE3) RIPL by the heat shock method and cultured overnight at 37°C on LB agar plates supplemented with 100 µg/mL ampicillin. *E. coli* BL-21 CodonPlus (DE3) RIPL with the target plasmid was added to fresh LB medium, supplemented with 100 µg/mL ampicillin. When the OD_600nm_ reached 0.4, the incubation temperature was decreased to 25°C and IPTG was added to each culture at a final concentration of 0.1 mM to induce protein production during the overnight incubation. Subsequently, cells were pelleted and lysed by adding lysozyme and benzonase nuclease (Sigma) and by passing through a French pressure cell press twice. Cell lysate was separated by centrifugation (13,300 × *g*) at 4°C for 30 min. The recombinant proteins in the soluble fraction were purified using Strep-Tactin resins (IBA lifesciences) according to the manufacturer’s instructions. The purified proteins were assessed by sodium dodecyl sulfate-polyacrylamide gel electrophoresis (SDS-PAGE). Protein concentrations were measured by Nanodrop 2000c spectrophotometer based on their extinction coefficients and molecular weights.

#### Steroid extraction

Microbial cultures incubated with cortisol or 11OHAD were subjected to the steroid extraction process. A volume of 300 µL of culture medium was transferred to the glass coated 2 mL deep well plates (Thermofisher Scientific). To each well, 800 µL of HPLC/LCMS-grade ethyl acetate was added and mixed thoroughly. The plates were centrifuged at 2000 rpm for 5 minutes to facilitate phase separation. The upper organic layer was carefully collected and transferred to a new deep well plate. The extraction was done twice. Ethyl acetate was dried up by evaporation in a chemical fume hood.

#### Human Sterolbiome Discovery High-throughput (HSDH) assay

To analyse the samples with the HSDH assay, 98 μL phosphate-buffered saline buffer with 100 μM cofactor (NADPH/NADP^+^) was added to each well containing the extracts. Then vortex for 2 min to resuspend the extracts. Samples were then transferred to a new Nunc™ MicroWell™ 96-Well, Nunclon Delta-Treated, Flat-Bottom Microplate. 2 μL enzyme was added with a final concentration of 0.5 μM, and the system was mixed thoroughly, and the kinetic reactions were measured immediately with a BioTek plate reader at 340 nm wavelength.

#### Bacterial metabolite analysis with Liquid chromatography-mass spectrometry (LC-MS)

Bacterial metabolite analysis was performed using LC-MS. All samples were analyzed on a Waters Aquity UPLC coupled with a Waters Synapt G2-Si ESI MS (Waters Corp., Milford, MA, USA) coupled with a Waters Acquity UPLC BEH C18 column (1.7 μm particle size, 2.1 mm x 50 mm). The injection volume is 1 µL, and the column temperature was maintained at 40°C during the analysis. For the elution buffer, mobile phase A and mobile phase B were used: mobile phase A contained 95% water, 5% acetonitrile, and 0.1% formic acid; mobile phase B contained 95% acetonitrile, 5% water, and 0.1% formic acid. The LC eluents were introduced into the mass spectrometer equipped with electrospray ionization (ESI) with a positive ion mode for steroid analysis. During the elution process, initially, mobile phase A was 100% for 0.5 min. Over the next 5.5 min, mobile phase B linearly increased, reaching 70% at 6 min. Then, mobile phase B increased to 100% in 1 min and was maintained for 1 min. Afterwards, a steep reversal to the initial conditions was done within 0.1 min, and the running condition was maintained until the end at 10 min. The flow rate was 0.5 mL/min. The following optimized conditions were used: capillary voltage of 3 kV, desolvation temperature of 500°C, cone voltage of 25 V, collision energy of 4 eV, collision gas helium, source temperature of 120°C, cone gas flow of 10 L/h, and desolvation gas flow of 800 L/h. The mass range was 50-2000 Da. Chromatographs and mass spectrometry data analysis were done with Mass Lynx v4.1 (Waters) software.

#### Whole genome sequencing

Urinary isolates were incubated at 37°C in anaerobic PYG broth. During log growth, cells in cultures were harvested by centrifugation (4,000 rpm) at 4°C for 15 min. High-molecular-weight (HMW) DNA extraction, library preparation and sequencing were conducted at the Roy J. Carver Biotechnology Center at the University of Illinois at Urbana-Champaign. Cell pellets were washed and resuspended with 200 µL 1×PBS (Corning, VA). Cell lysis was done with 5 µL Metapolyzyme (Sigma) and a 2-hour incubation at 37°C. Additional lysis was done on all samples with the MagAttract HMW DNA Kit (Qiagen, MD). Chloroform was added, and samples were centrifuged at 10,000 × g for 2 min to isolate the supernatant containing the DNA. DNA purification was subsequently carried out following the MagMAX™ Plant DNA Isolation Kit (Thermo Fisher Scientific, MA), with buffers prepared as per the manufacturer’s instructions. DNA was eluted in 50 µL of elution buffer and quantified using Qubit (Thermo Fisher Scientific) and fragment sizes were evaluated in the Femto Pulse system (Agilent, CA). The HMW DNA was sheared with a Megaruptor 3 to an average fragment length of 10 kb. Sheared DNA was converted to a library with the SMRTBell Express Template Prep kit 3.0. The library was sequenced on 1 SMRTcell 8M on a PacBio Revio using the CCS sequencing mode and a 30-hour movie time. CCS analysis was done in an instrument with SMRTLink V13.1 using the following parameters: ccs --min-passes 3 --min-rq 0.99.

#### Whole genome sequencing analysis

FastQC v0.11.8^63^ was used to evaluate read quality. SeqKit v2.0.0^64^ was used to calculate the statistics of the sequencing files for each microbial isolate. Read number, sum of the read length, minimum read length, average read length and maximum read length were shown in **Supplementary Table 7.** Flye v2.9^65^ was used to assembly the reads (enough for 50 fold coverage) chosen using the parameters: --asm-coverage. Assembly quality and completeness were evaluated using QUAST v5.0.2^66^ and BUSCO v5.5.0^67^ respectively. Annotations were performed using Prokka v 1.14.6^68^. CGView Server was used to make the circular genome maps^69^. GTDB-Tk v2.3.0 (database R220)^70^ was used for the taxonomy assignment.

#### Pangenome analysis of *P. lymphophilum* strains

The sequenced genomes of 18 strains of *P. lymphophilum* that were isolated in this study and 4 genomes that were downloaded from NCBI were included for the pangenome analysis (**Supplementary Table 2-3**). All of the genomes were evaluated using QUAST v5.0.2^66^ and BUSCO v5.5.0^67^ respectively. Annotations were performed using Prokka v 1.14.6^68^. The files generated were used for the pangenome analysis. The *P. lymphophylum* pangenome was performed using the Roary tool v3.11.2^71^. Subsequently, FastTree v2.1.11^72^ and a Roary script (roary_plots.py) was used to visualize a presence-absence matrix of core and accessory genes shared between genomes. Heaps Law’s alpha parameter (to estimate whether the pangenome is closed or open) and core genome predicted size were estimated using the micropan R package^73^. HMMER v3.3.1 was used to search the proteins DesA, DesB, DesE, and DesG based on the experimentally verified sequences. Pyani v0.2.10 was used to calculate the average nucleotide identity (ANI) with -m ANIm^74^. UpSetR package was used to visualize the unique and intersection genes for each *P. lymphophilum* genome^75^. EggNOG-mapper v.2 online tools^76^ were used for the protein functional annotations based on the Clusters of Orthologous Genes (COG) database. KOALA v.3.1 (KEGG Orthology And Links Annotation)^77^ was used to study the metabolic pathways with the Kyoto Encyclopedia of Genes and Genomes (KEGG) database. The COG and KEGG categories were calculated for the set of core genes, accessory genes, and unique genes.

#### Scanning electron microscopy (SEM)

*A. massiliense* CFH19 strain was cultured in PYG broth in an anaerobic chamber. Karnovsky’s fixative, containing 2% glutaraldehyde and 2.5% paraformaldehyde was used to fix the bacterial isolates. Cultures (500 µL) in log phase were mixed with fixative (500 µL), vortexed, and then centrifuged 4,000 x *g* for 5 min at 4°C. Supernatant was discarded, and the pellet was washed 3 times with fixative (500 µL) and then stored at 4°C. Samples were sent to Materials Research Laboratory Central Research Facilities (University of Illinois at Urbana-Champaign, Urbana, Illinois, United States) for SEM imaging.

#### Whole cell steroid conversion assay by *A. massiliense* CFH19

Frozen stock of *A. massiliense* CFH19 strain was revived in PYG medium in anaerobic condition. For the isomerase pathway exploration, log phase CFH19 culture was incubated with 50 uM DHEA, androstenediol, and androstenedione in PYG medium. 500 μL aliquots were collected at different time points (0, 24, 48 and 72 hours). These samples were centrifuged in a microcentrifuge to pellet the bacterial cells, and the supernatants were utilized for steroid quantification.

#### Gene activities identification by heterologous expression

The target inserts in the pET-IDT C His (**Supplementary Table 6**) were synthesized by Integrated DNA Technologies (IDT, Coralville, IA, USA). Recombinant plasmids were transformed into *E. coli* BL21-CodonPlus(DE3)-RIPL by heat shock method at 42°C for 30 seconds and cultured overnight at 37°C on LB agar plates supplemented with 50 µg/mL kanamycin. Selected colonies were transferred into the fresh LB broth with 50 µg/mL kanamycin for 24 h at 37°C. Cells were harvested by centrifugation (4,000 × *g*) at 4°C for 20 min. The plasmids were extracted using the QIAprep Spin Miniprep kit (Qiagen, Valencia, CA, USA). The extracted plasmids were sent to Plasmidsaurus, Inc. (Eugene, OR, USA) for whole plasmid sequencing using Oxford Nanopore Technology with custom analysis and annotation. After the verification, the *E. coli* BL-21 CodonPlus (DE3) with the target inserts were incubated in the LB broth in the air at 37°C. When the OD_600nm_ reached 0.4, the culture was transferred to the Huagate tubes, and the headspace was replaced with N_2_. IPTG was added to each culture at a final concentration of 0.1 mM, and substrates were added to each culture at a final concentration of 50 μM. The incubation temperature was decreased to 25°C. After 24 hour incubation. Culture was centrifuged (4,000 × *g*) at 4°C for 20 min to separate the cells and the supernatant. The supernatant was used for the metabolite analysis. Two parts ethyl acetate and one part bacterial culture supernatant were thoroughly mixed by vortexing for 1 min. Next, the ethyl acetate layer was carefully collected and transferred to new tubes. The extraction process was repeated, and the collected top layers were evaporated with nitrogen gas and dissolved in 200 µL LC-MS grade methanol. For samples where substrates and end products were to be quantified, an internal standard mixture of androstenedione-D7 and testosterone-D3 was added before extraction. The samples were analysed by LC-MS, and the concentrations of DHEA, androstenedione, androstenediol and testosterone were normalized based on the under-curve area of the internal standards accordingly.

#### Phylogenetics of DirA and DirB

The DirA and DirB protein sequences of *A. massiliense* CFH19 were used as queries for BLASTP searches against the NCBI’s NR protein database. The protein sequences from the top 100 results were aligned using MUSCLE v. 5.1 with default parameters^78^. Ambiguously aligned positions were removed with trimAl^79^. Each phylogeny was inferred by IQ-TREE v2.3.5^80^ with one thousand ultrafast bootstrap pseudoreplicates and one thousand SH-like approximate likelihood-ration test replicates, and -m MFP parameter was included for the auto model selection^81^. An online tool (iTOL) was used for the tree display and annotation^82^.

#### Computational Structural Biology

Structural models of the enzymes DirA and DirB were generated with AlphaFold3^39^. Modelling was performed with NAD⁺ or NADH included in order to stabilize the catalytic cleft during structure prediction. The resulting structures were protonated at physiological pH and visually inspected in VMD^83^. All subsequent computational analyses used these structures without modification.

Sterol ligands, including DHEA, androstenediol, androstenedione, and testosterone, were downloaded from PubChem in mol2 format^84^. Each ligand was protonated at pH 7.4 and energy-minimized using OpenBabel, then converted to pdbqt format for docking simulations. Protein receptors were similarly processed with OpenBabel to assign atom types and prepare pdbqt files. Docking simulations were performed using AutoDock Vina^85^. Search grids were centered on the predicted catalytic pocket to permit sampling of both the 3′ and the 17′ orientations required for the proposed reaction sequence. For each enzyme-ligand pair, the lowest-energy poses were inspected manually in VMD. Poses were evaluated based on binding energy, alignment relative to NAD⁺ or NADH, and the spatial relationship between the ligand and the catalytic residues.

Molecular dynamics simulations were performed with NAMD 3 using the QwikMD interface. Each protein-ligand complex was solvated in a TIP3P water box with at least 12 Å padding and ionized to 150 mM NaCl. Systems underwent 1000 steps of energy minimization, followed by gradual heating from 10 K to 300 K in increments of 1 K under NVT conditions. Equilibration was performed for 0.5 ns at 300 K and 1 atm, followed by 10 ns of unrestrained production dynamics under identical conditions. Trajectories were analyzed in VMD to monitor ligand stability, orientation relative to the catalytic residues, and the position of reactive groups with respect to the NAD⁺ or NADH cofactor.

#### Data analysis

Data analysis was performed with R version 4.3.0^86^. Data are shown as means ± standard deviations (SD) when data are normalized. Data are shown in median and interquartile ranges when skewed.

#### Data availability

The raw genome sequencing data and assemblies are deposited at the NCBI with accession numbers PRJNA1372755 and PRJNA1372814, respectively.

## Acknowledgements

J.M.R. would like to express gratitude to both the Cancer Center at Illinois and the Center for Advanced Study at Illinois for the financial support and protected time to pursue this work. We would like to thank Alvaro Hernandez, Director of DNA Services Facility at the Roy J. Carver Biotechnology Center at Urbana-Champaign and Christopher J. Fields, Director of High-Performance Biology Computing at the Roy J. Carver Biotechnology Center for nucleic acid sequencing and bioinformatic assistance. We thank Kristen M. Flatt for SEM images of bacterial isolates. We thank Furong Sun, Director of Mass Spectrometry Laboratory at Urbana-Champaign for LC-MS analysis. All chemical structures were created with ChemdDoodle. This work was supported by grants from the National Institutes of Health (R01 GM145920 [J.M.R]), Cancer Center at Illinois Seed Grant [J.M.R, J.I., J.E. Jr, H.R.G.], UIUC Department of Animal Sciences Matchstick grant, and Hatch ILLU-538-916. F.F. was supported through a Fulbright Fellowship. B.B. was supported by NIGMS Diversity Supplement on R01 GM134423. Computational work supported by the National Science Foundation grant MCB-2143787 and NIH R24 GM145965 [R.C.B.]. The authors acknowledges financial support from the Alabama Graduate Research Scholars Program (GRSP) funded through the Alabama Commission for Higher Education and administered by the Alabama EPSCoR.

## Author contributions

J.M.R. conceptualized the HSDH assay, identified *dir* gene candidates, and conceptualized and supervised the overall study and wrote the first draft and edited subsequent drafts. J.I., supervised culturomics, and contributed to the second and subsequent drafts. R.C.B. supervised and performed the computational structural biology work, and contributed to the first draft. R.S.L.R performed computational structural biology and developed catalytic models. J.W.E Jr., S.L.D., H.R.G., provided guidance and supervision of key aspects of the study. T.W. conceptualized the study, wrote the draft, developed the HSDH assay, developed the method for LC-MS analysis, performed the whole genome sequencing analysis, pangenome analysis and phylogenetic analysis, characterized the *dir* genes and completed the pathways. S.A. processed patient samples, urine culturomics, HSDH assay plate preparation and LC/MS analysis. J.O.I., D.C., and V. P. performed urine culturomics and LC/MS analysis. B.B. performed heterologous expression and LC/MS. F.F. performed bioinformatic analysis. M.A.B and E.T. recruited patients. P.M., D.D., D.O., M.A.B., and E.T. collected patient urine samples. J.M.R., J.I., T.W., and S.A. analysed data and wrote the manuscript with input from all the co-authors.

## Extended data figures

**Extended Data Figure 1:**
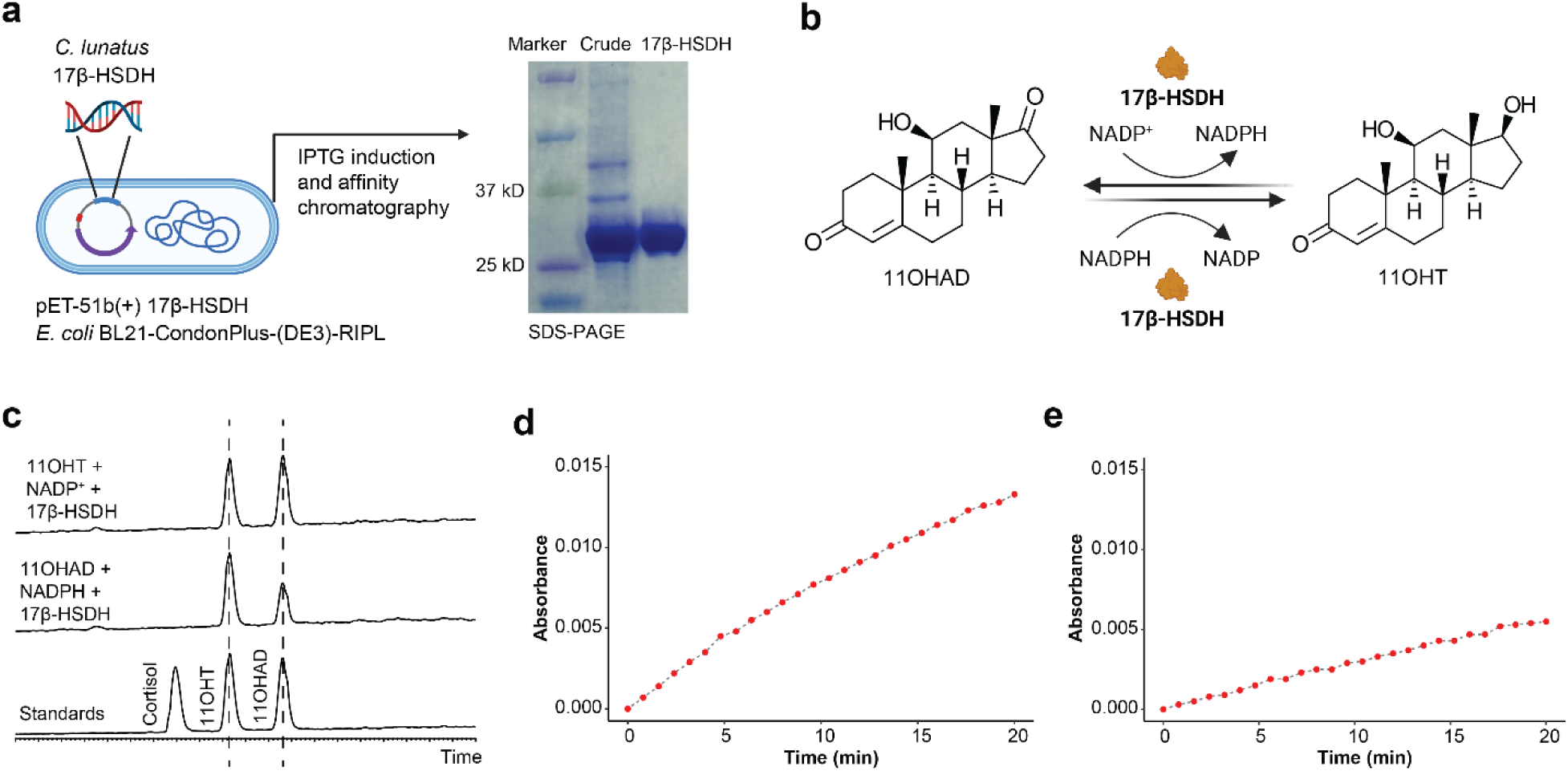
Characterisation of the 17β-HSDH from *Cochliobolus lunatus*. **a,** The cDNA encoding 17β-HSDH was cloned into pET-51b(+) plasmid and expressed in *E. coli* BL21-CondonPlus-(DE3)-RIPL. **b,** 17β-HSDH from *C. lunatus* converted 11OHAD to 11OHT using NADPH as cofactor and converted 11OHT to 11OHAD using NADP^+^ as cofactor. **c,** LC-MS chromatograms confirmed the 17β-HSDH enzymatic activities. **d,** The change of the optimal density (absorbance) using NADPH as cofactor overtime at 340 nm when 11OHAD was converted to 11OHT by 17β-HSDH from *C. lunatus*. **e,** The change of the optimal density (absorbance) using NADP^+^ as cofactor overtime at 340 nm when 11OHT was converted to 11OHAD by 17β-HSDH from *C. lunatus*.

**Extended Data Figure 2:**
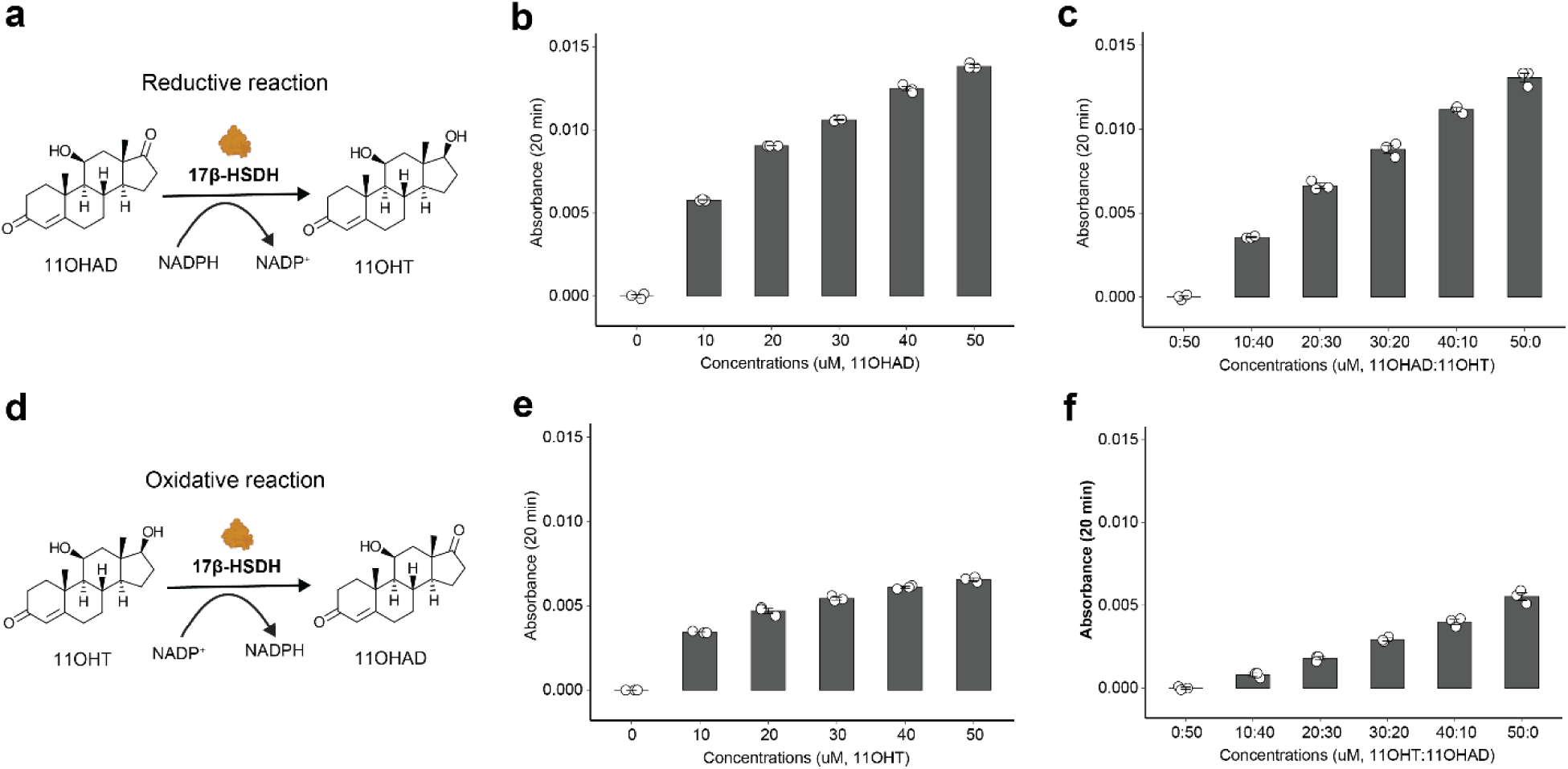
Effect of the substrate concentrations on HSDH assay absorbance at 340 nm. **a,** 11OHAD was converted to 11OHT by 17β-HSDH from *C. lunatus*, concomitantly, NADPH was converted to NADP^+^. **b,** The change of the optimal density (absorbance) increased with increasing 11OHAD concentrations. **c,** The total concentration of 11OHAD and 11OHT is 50 µM. The concentrations showed the ratio of 11OHAD and 11OHT. The change of the optimal density (absorbance) increased with increasing 11OHAD concentrations. **d,** 11OHT was converted to 11OHAD by 17β-HSDH from *C. lunatus*, concomitantly, NADP^+^ was converted to NADP^H^. **e,** The change of the optimal density (absorbance) increased with increasing 11OHT concentrations. **f,** The total concentration of 11OHT and 11OHAD is 50 µM. The concentrations showed the ratio of 11OHT and 11OHAD. The change of the optimal density (absorbance) increased with increasing 11OHT concentrations.

**Extended Data Figure 3:**
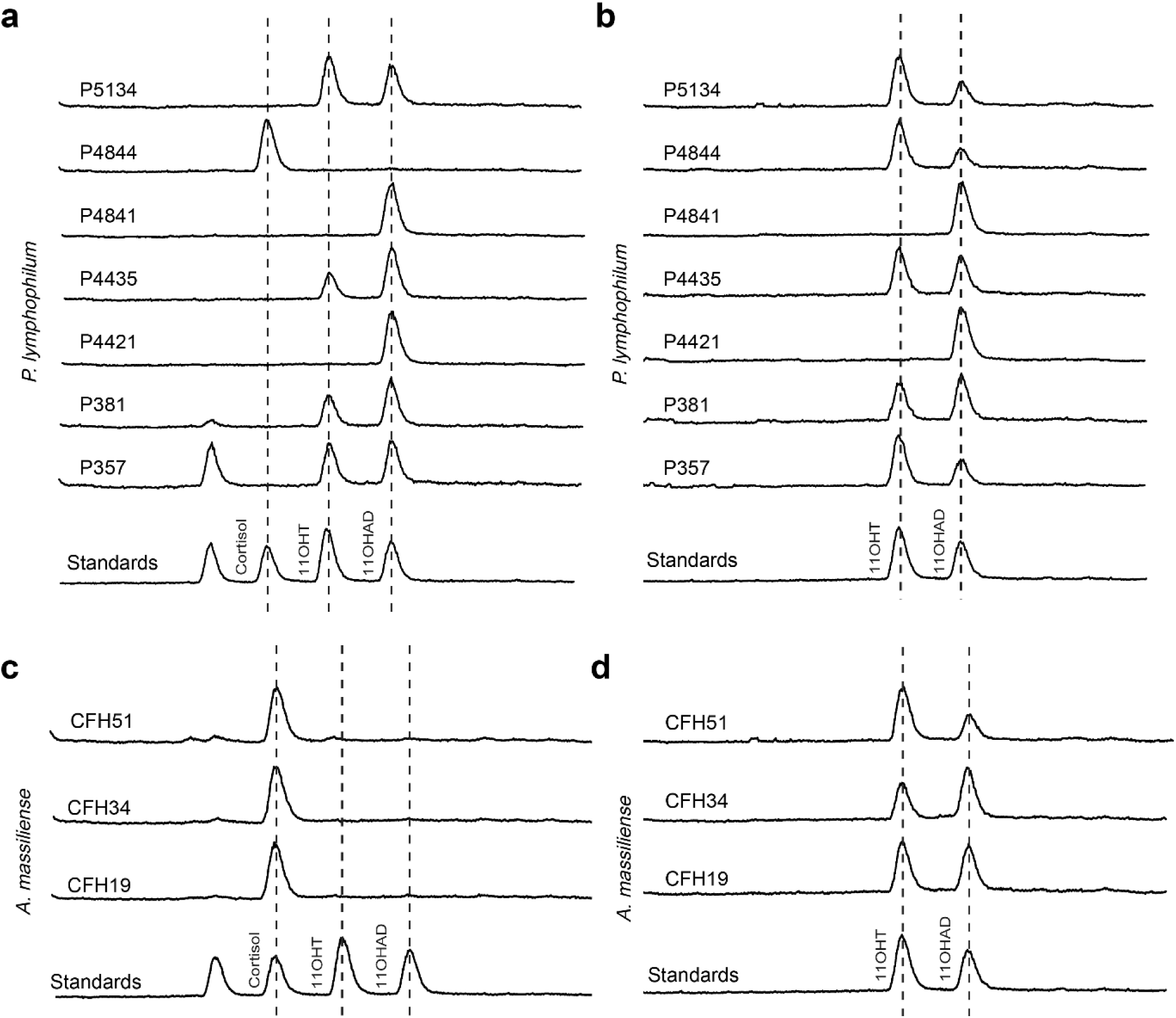
LC-MS chromatograms of *Propionimicrobium lymphophilum* and *Actinobaculum massiliense* strains using cortisol (a, c) or 11OHAD (b, d) as substrates. *P. lymphophilum* strain C741 cannot grow after isolation, and the LC-MS chromatograms of this strain are not included.

**Extended Data Figure 4:**
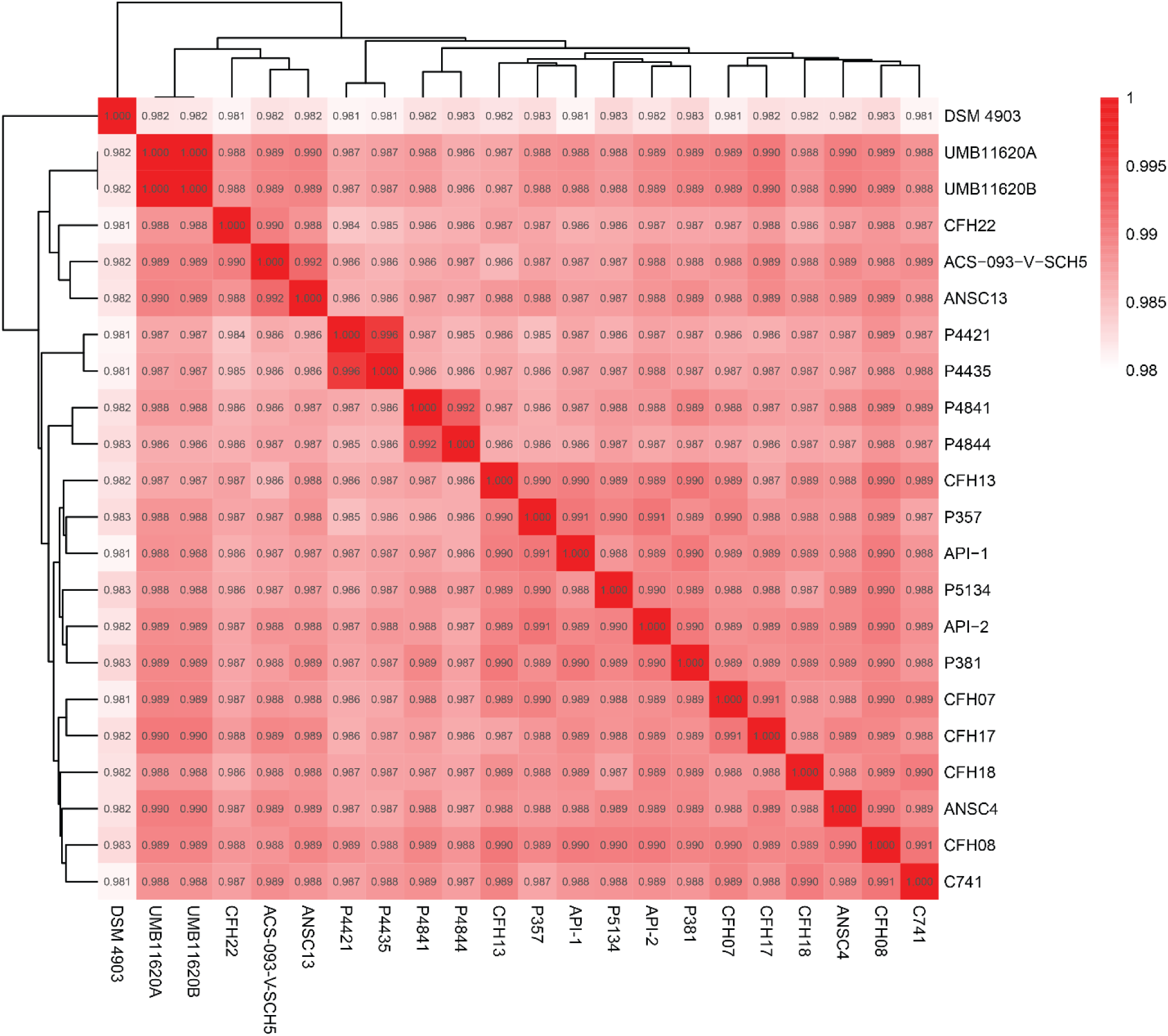
Average nucleotide identity between *Propionimicrobium lymphophilum* strains.

**Extended Data Figure 5:**
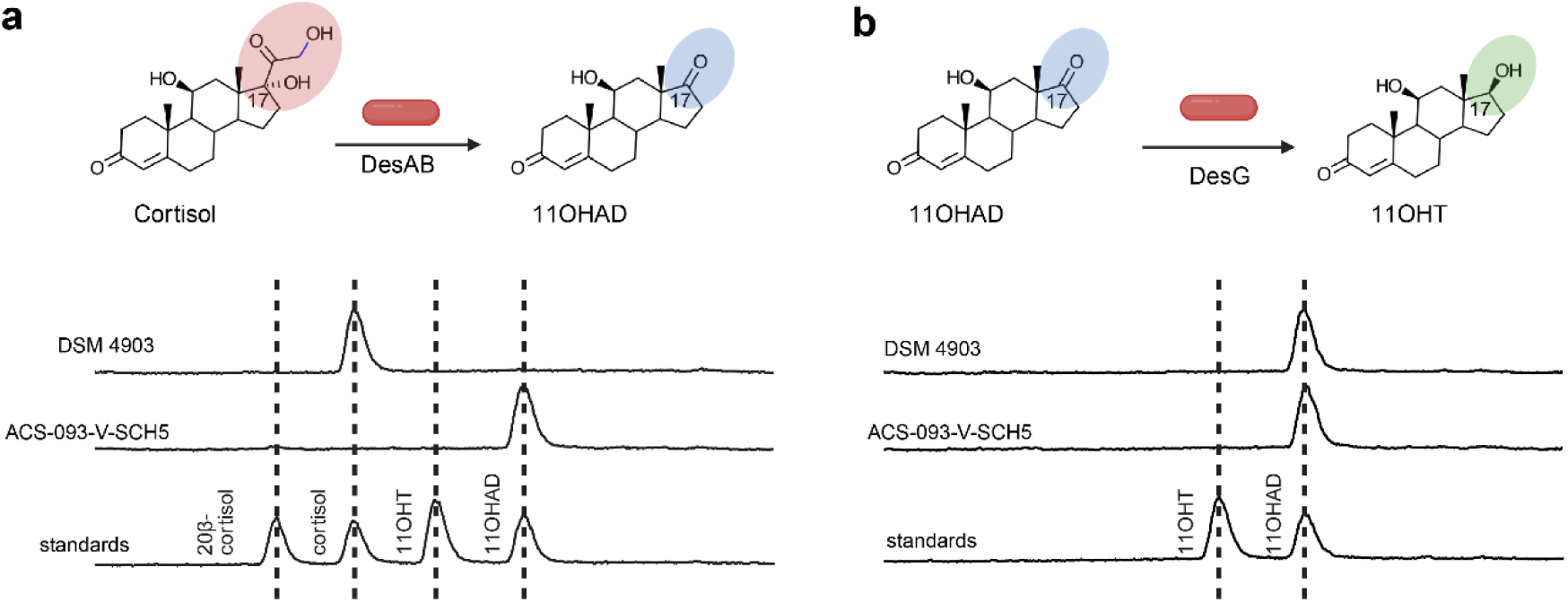
LC-MS chromatograms to show the (a) DesAB activities and (b) DesG activities of *Propionimicrobium lymphophilum* DSM 4903 and ACS-093-V-SCH5.

**Extended Data Figure 6:**
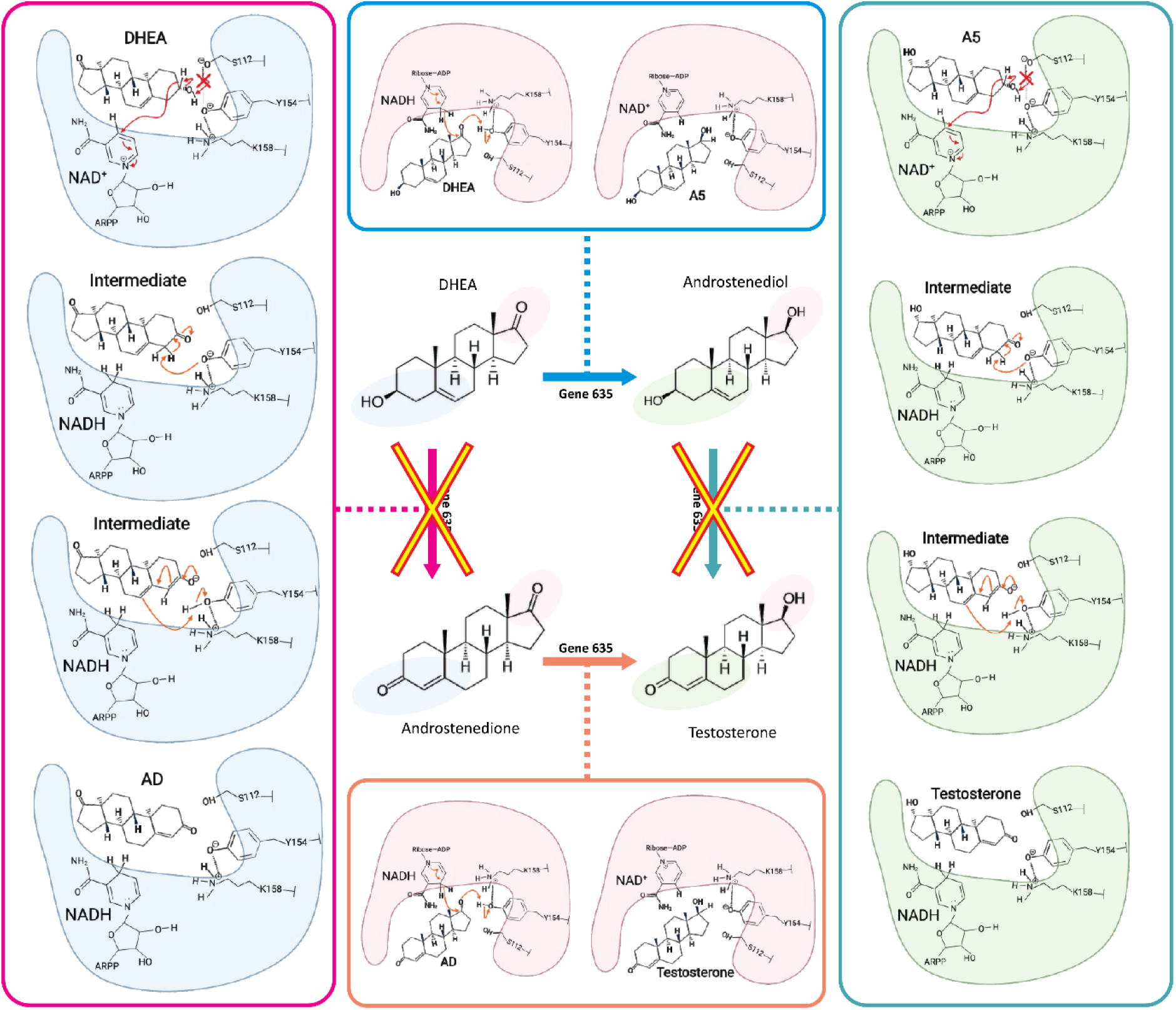
Proposed catalytic mechanism of DirB (Gene 635) in steroid conversion. Schematic representation of the reaction pathways catalyzed by DirB. The pink pathway (left) illustrates the inability to convert DHEA to androstenedione (AD). The orange pathway (bottom) depicts the reduction of androstenedione to testosterone based on productive binding of the 17′ side facing the active pocket and proton donation by NADH. The blue pathway (top) represents the conversion of DHEA to androstenediol (A5), while the cyan pathway (right) shows the inability to convert androstenediol to testosterone.

**Extended Data Figure 7:**
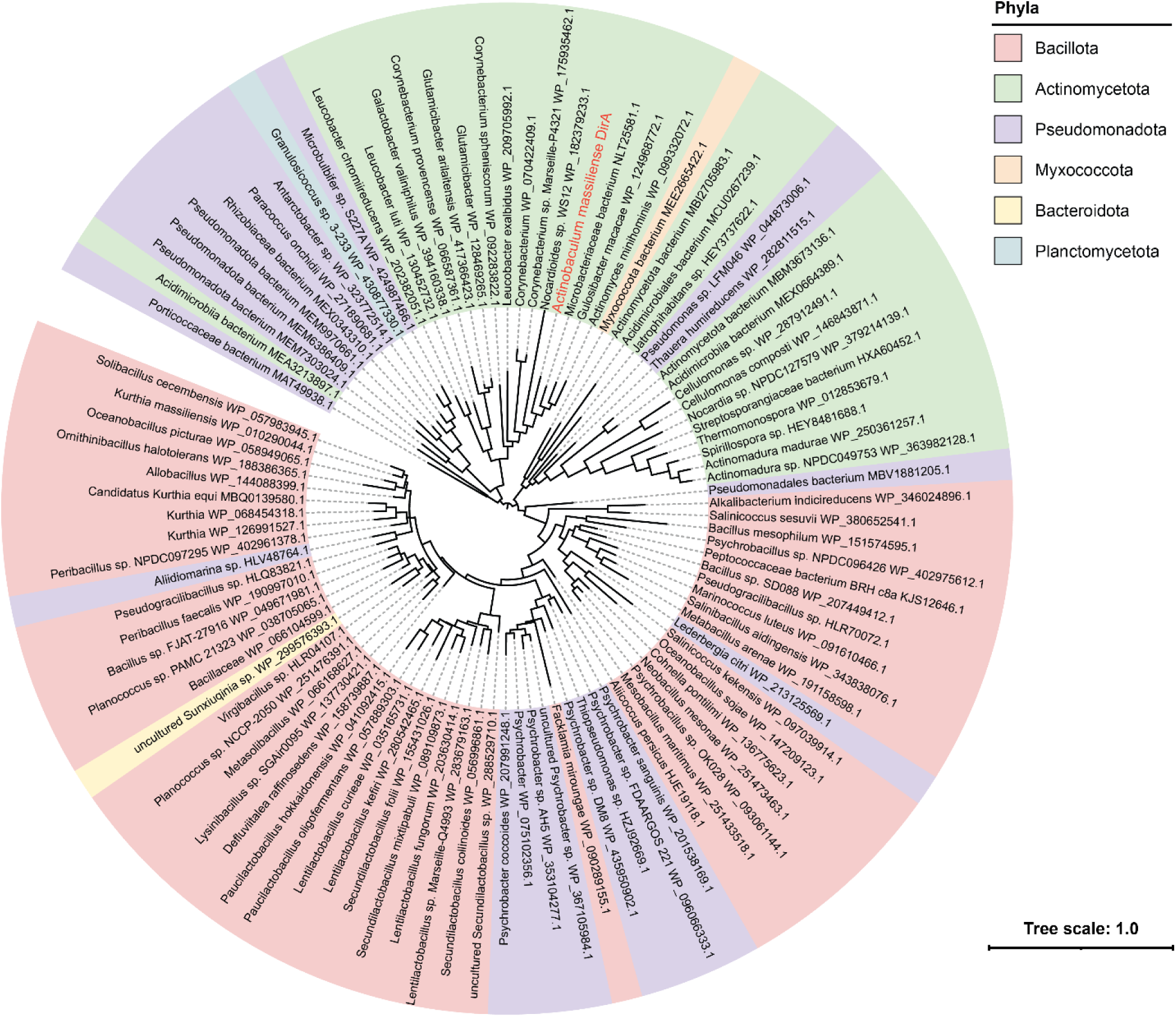
Maximum likelihood phylogenetic tree of DirA. The tree is colored by taxonomic affiliation as displayed in the legend.

**Extended Data Figure 8:**
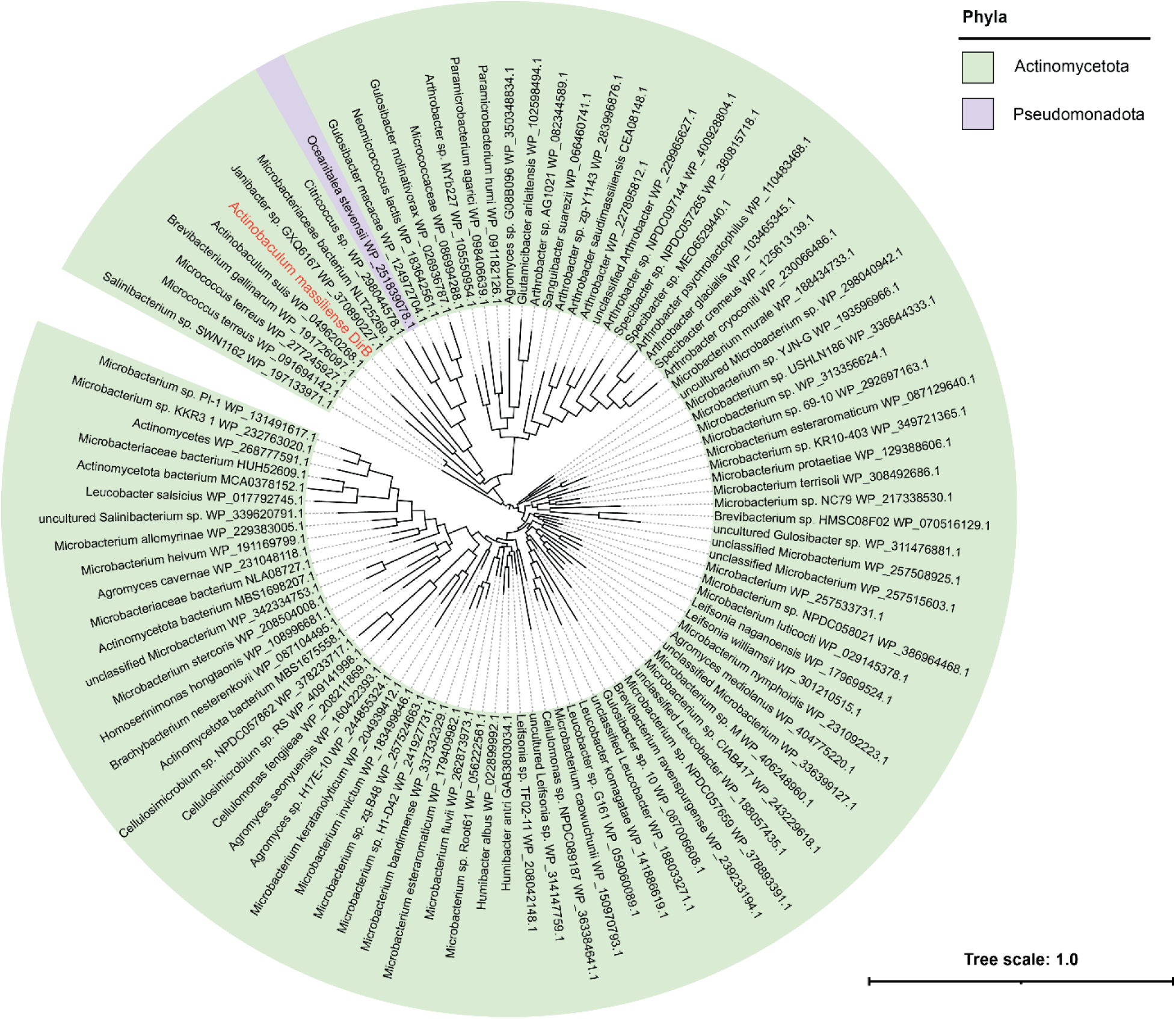
Maximum likelihood phylogenetic tree of DirB. The tree is colored by taxonomic affiliation as displayed in the legend.

**Extended Data Figure 9:**
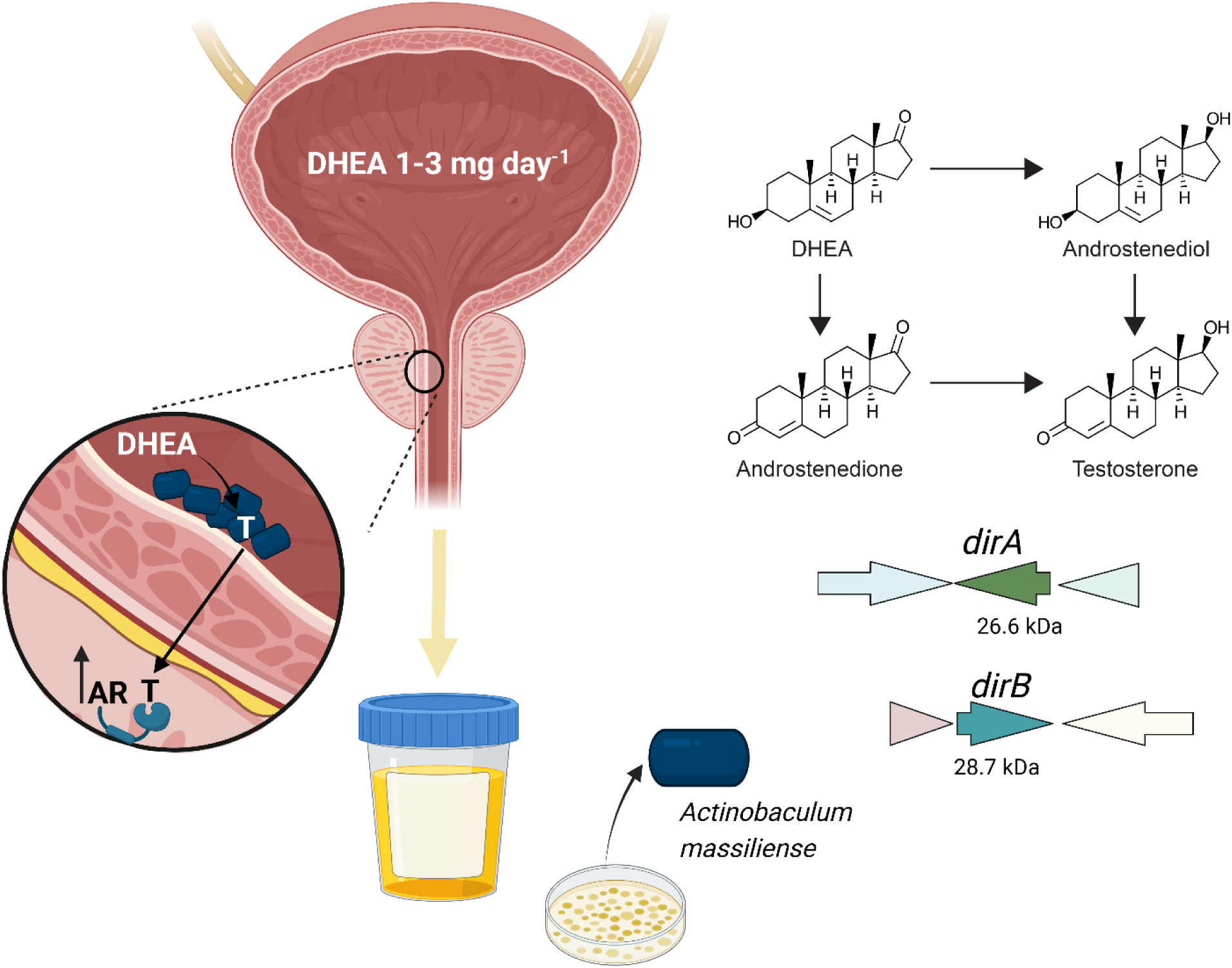
Schematic representation of the findings of this study as well as the hypothesis that androgen producing taxa convert DHEA to testosterone (T) in the prostatic urethra, along testosterone to passively diffuse into prostate tissue where it actives the androgen receptor (AR).

