## Supplementary Information for "The urinary pathobiont *Actinobaculum massiliense* generates androgens via the *dirAB* pathway"

Short title: Androgen biosynthesis via the *dirAB* pathway

### Supplementary files

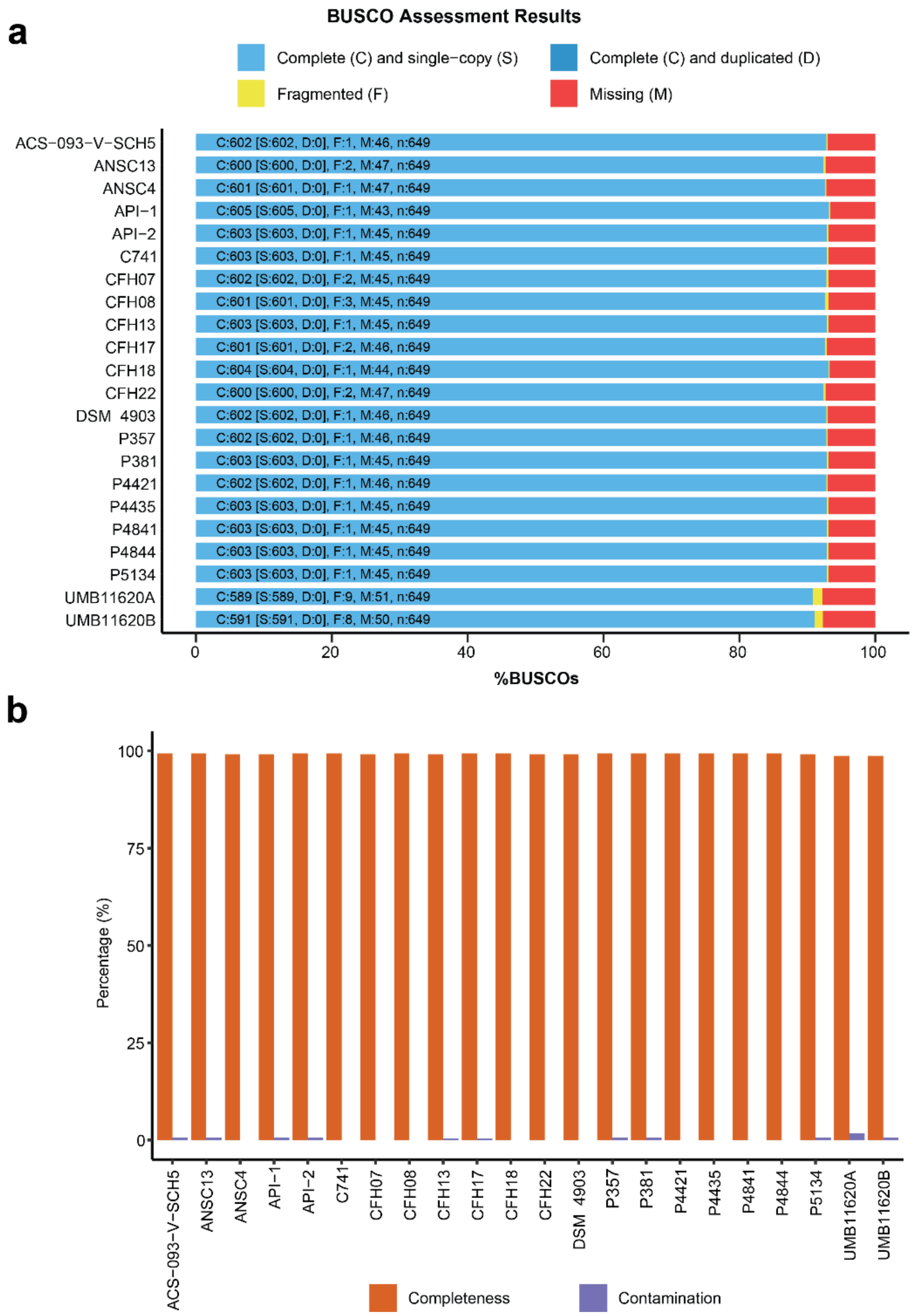


**Supplementary Figure 1: Genome quality of the *P. lymphophilum* strains assessed by BUSCO (a) and CheckM (b).**

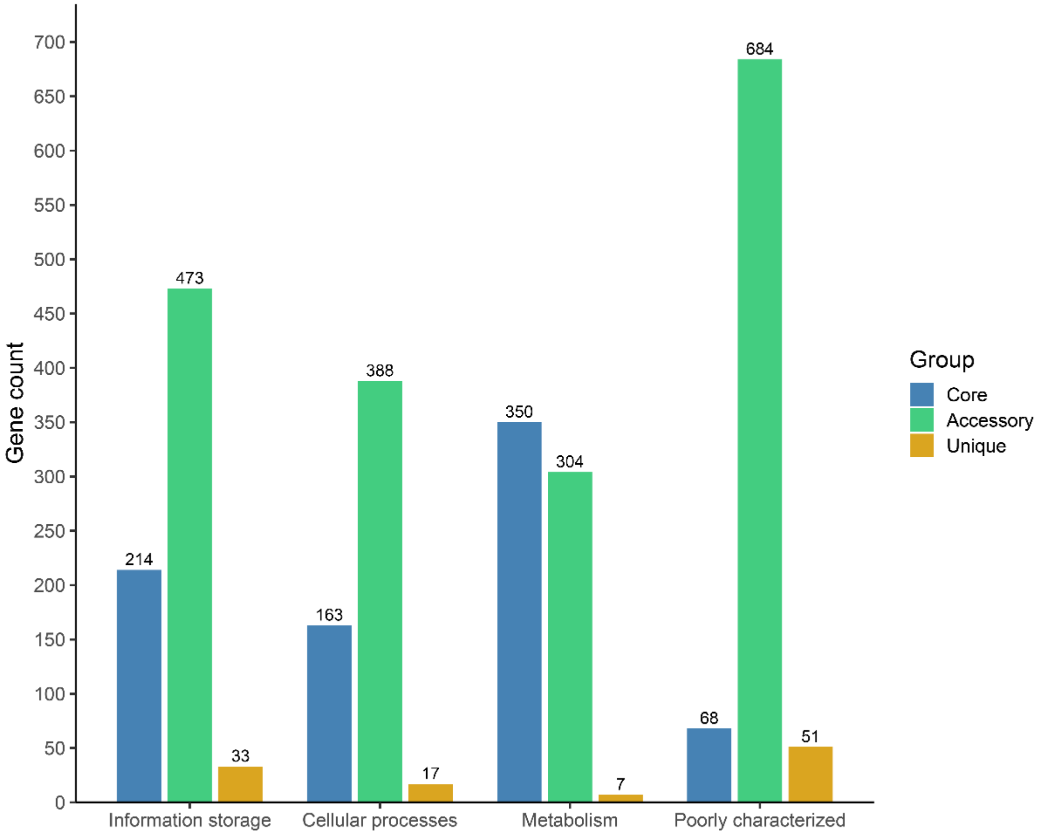


**Supplementary Figure 2: Clusters of Orthologous Genes (COG) analysis of *P. lymphophilum* pangenome. Total genes involved in Information Storage and Processing, Cellular Processes and Signaling, Metabolic Functions, or poorly characterized. Corresponding COG code and gene count of each category are listed in Supplementary Table 4.**

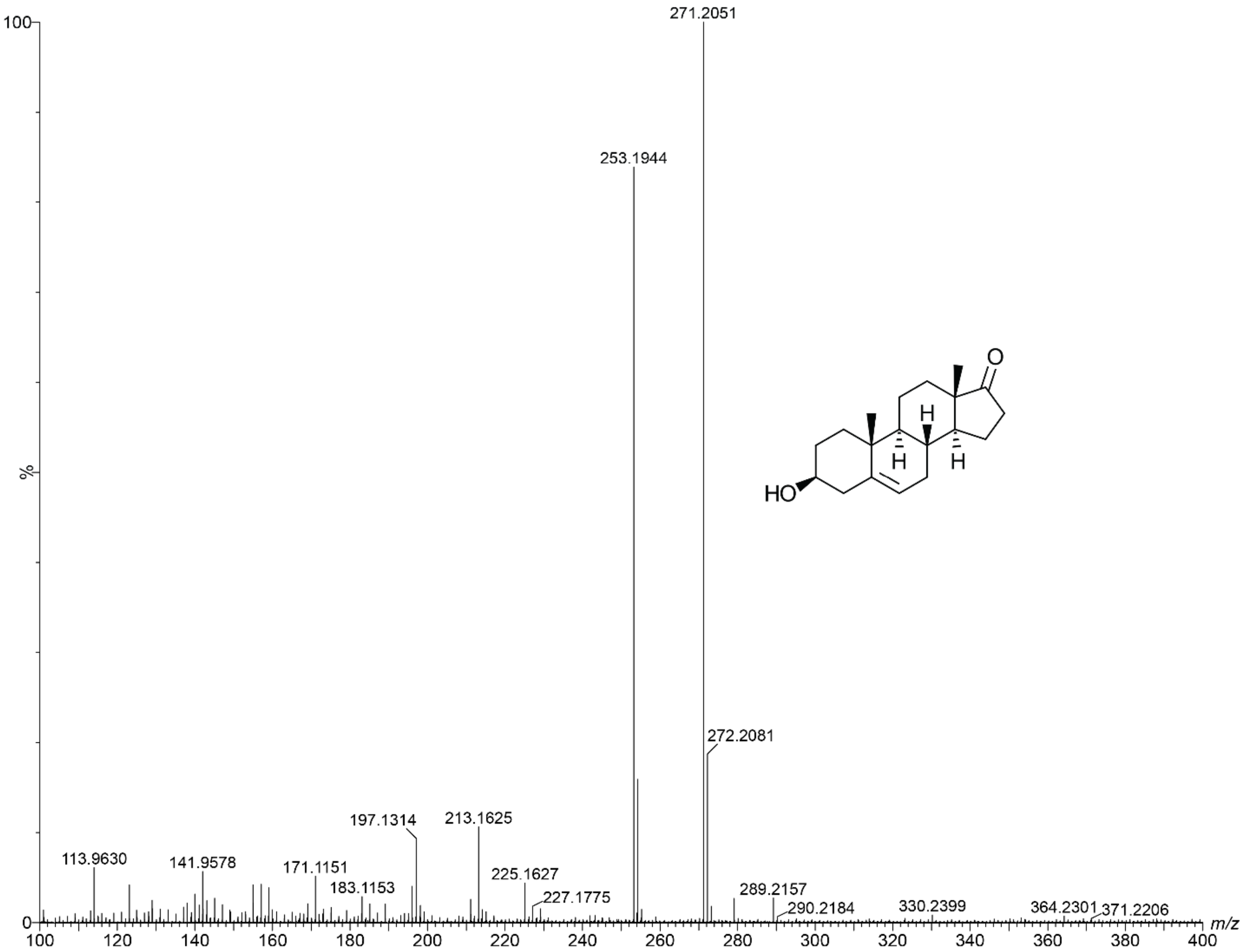


**DHEA**

**Supplementary Figure 3: High-resolution mass spectrum of DHEA in the positive mode. *m/z* 271.2051 was used for the quantification analysis in this study.**

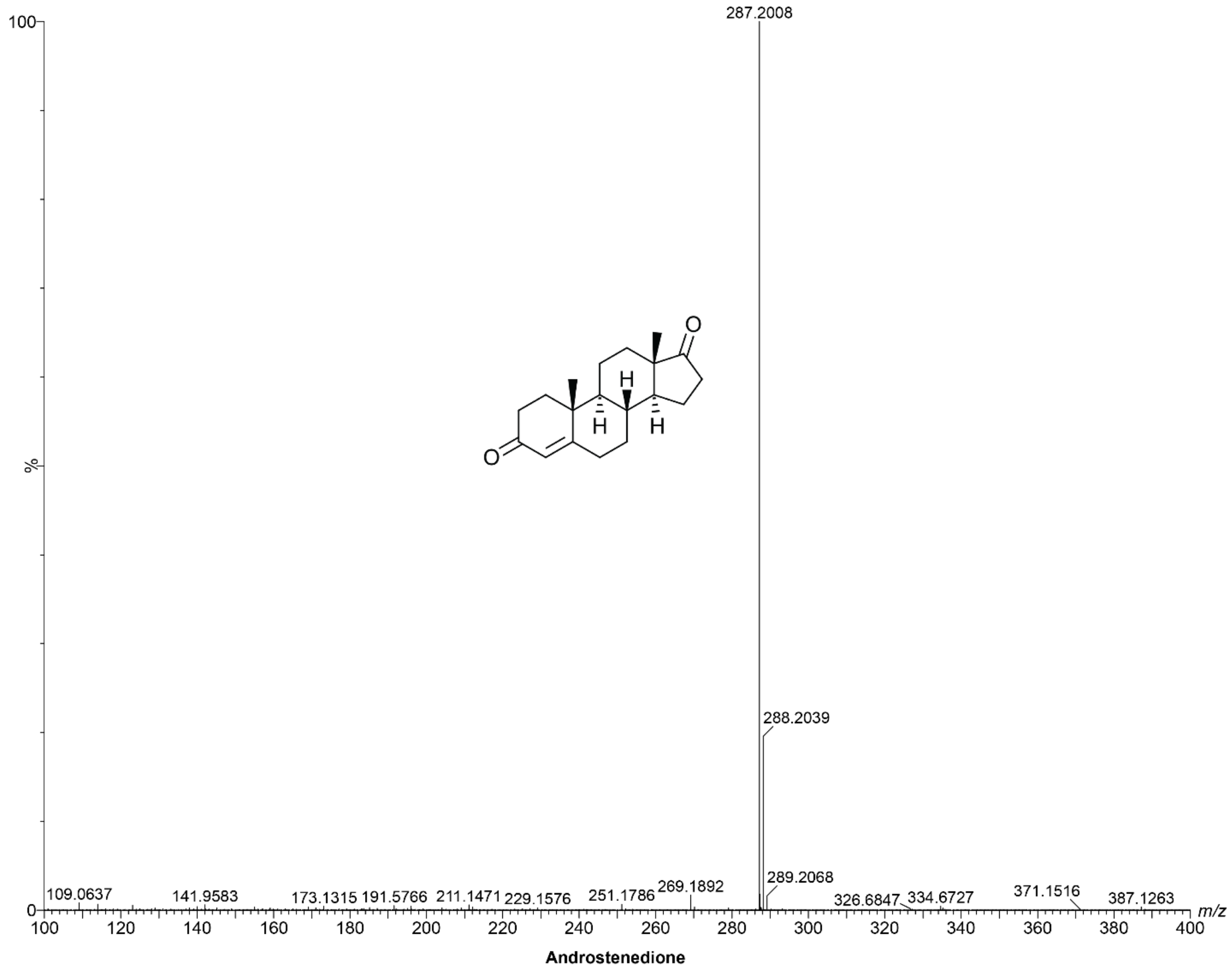


**Supplementary Figure 4: High-resolution mass spectrum of androstenedione in the positive mode. *m/z* 287.2008 was used for the quantification analysis in this study.**

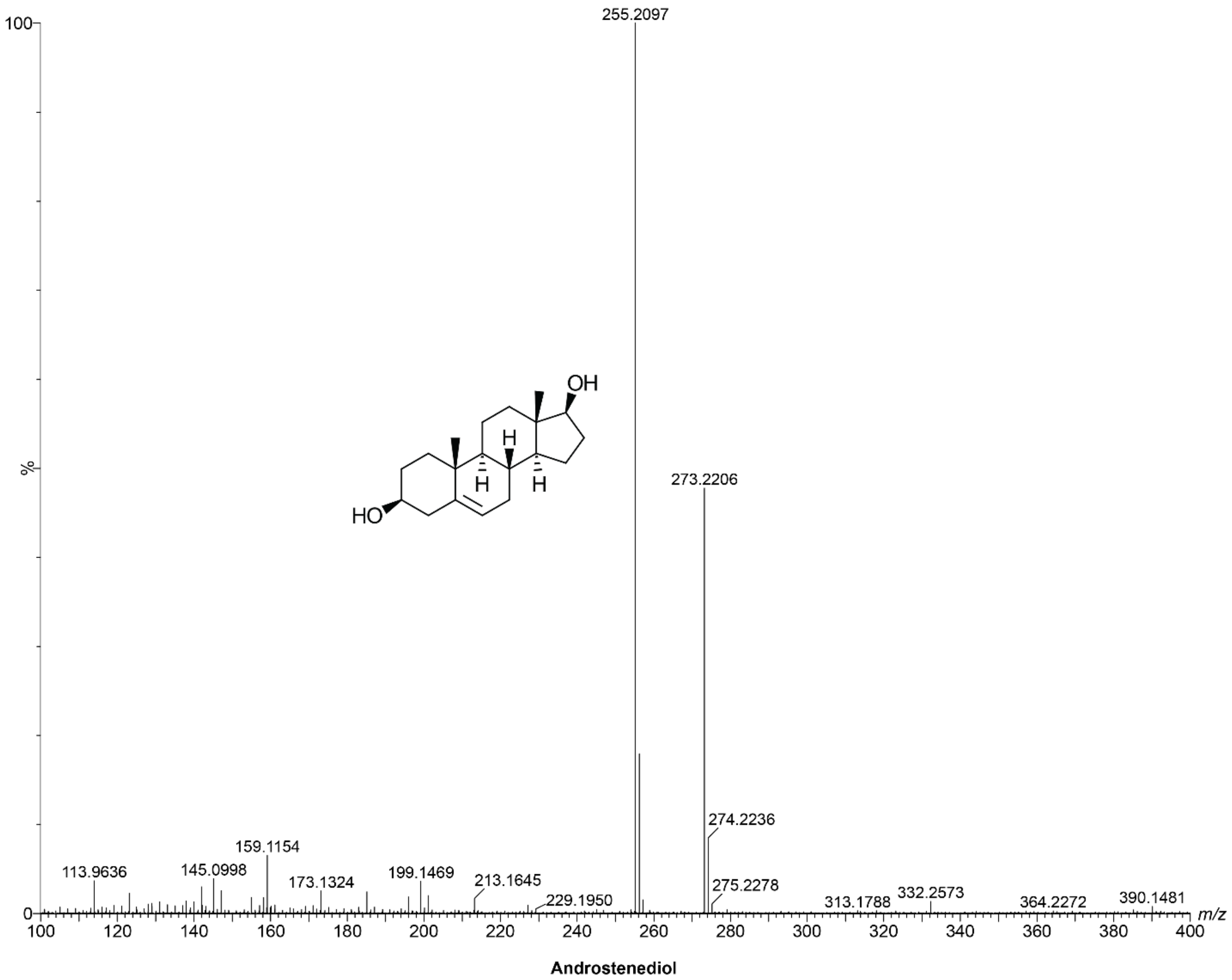


**Supplementary Figure 5: High-resolution mass spectrum of androstenediol in the positive mode. *m/z* 255.2097 was used for the quantification analysis in this study.**

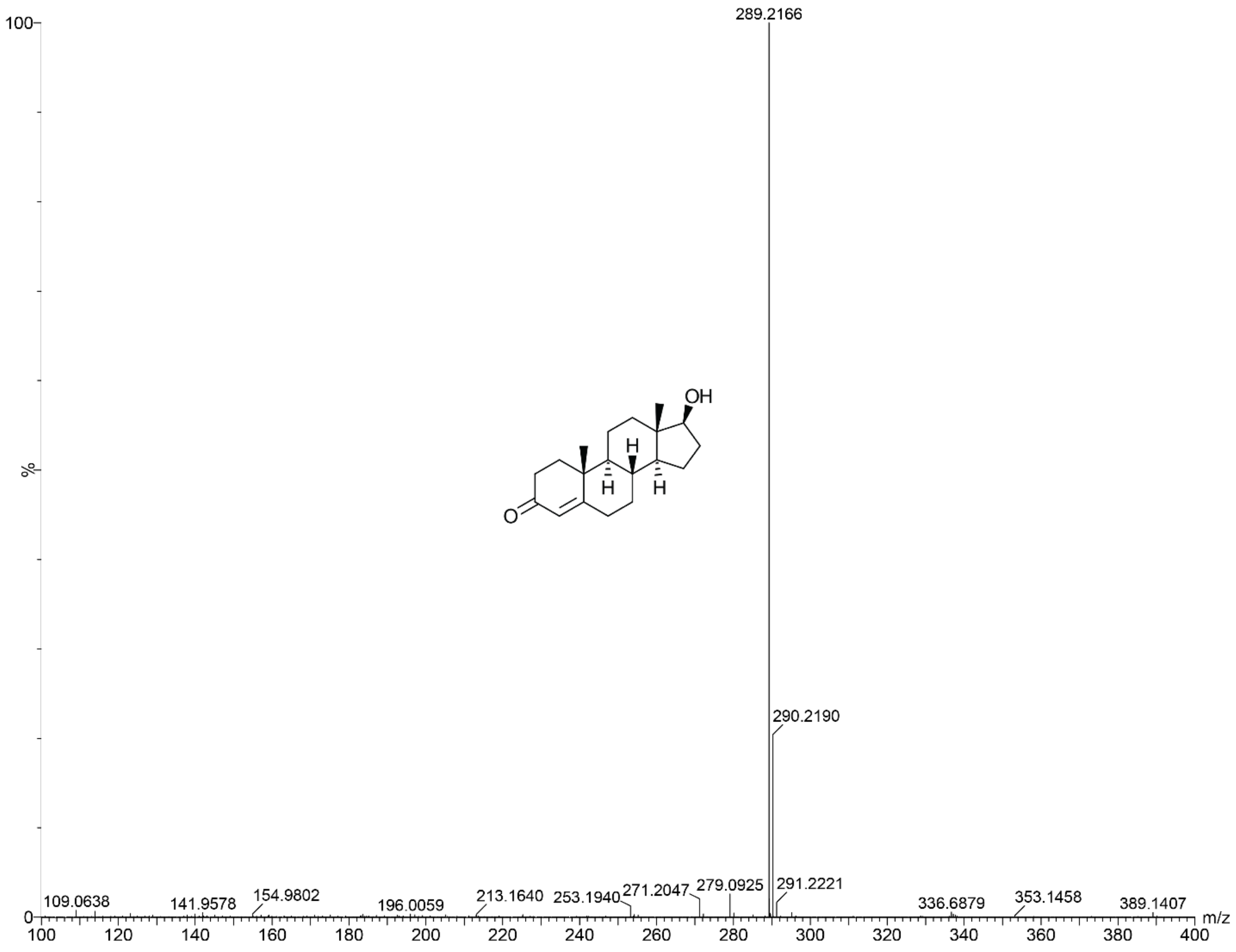


**Testosterone**

**Supplementary Figure 6: High-resolution mass spectrum of testosterone in the positive mode. *m/z* 289.2166 was used for the quantification analysis in this study.**
